# Understanding the photophysics and structural organization of photosynthetic proteins using model lipid membranes assembled from natural plant thylakoids

**DOI:** 10.1101/2020.09.15.296665

**Authors:** Sophie A. Meredith, Takuro Yoneda, Ashley M. Hancock, Simon D. Connell, Stephen D. Evans, Kenichi Morigaki, Peter G. Adams

## Abstract

The light-harvesting (LH) biomembranes from photosynthetic organisms perform solar energy absorption and transfer with high efficiency. There is great interest in the nanoscale biophysics of photosynthesis, however, natural membranes are complex and highly curved so can be challenging to study. Here we present model photosynthetic “hybrid membranes” assembled from a combination of natural LH membranes and synthetic lipids deposited into a patterned polymerized lipid template on glass. This arrangement offers many advantages over previous model systems including: a sufficiently complex mixture of natural proteins to mimic the biological processes, a modular self-assembly mechanism, and a stabilizing template promoting the formation of supported lipid bilayers from complex natural membranes with high protein content (that would not otherwise form). These hybrid membranes can be used as a platform to delineate the complex relationship between LH energy pathways and membrane organization. Atomic force microscopy and fluorescence lifetime microscopy revealed that hybrid membranes have an elongated fluorescence lifetime (∼4 ns) compared to native membranes (∼0.5 ns), a direct consequence of reduced protein density and an uncoupling of protein-protein interactions. We observed the real time self-assembly and migration of LH proteins from natural membrane extracts into the hybrid membranes and monitored the photophysical state of the membranes at each stage. Finally, experiments utilizing our hybrid membranes suggest that assays currently used in the photosynthesis community to test the electron transfer activity of Photosystem II may have non-specific interactions with other proteins, implying that new methods are needed for reliable quantification of electron transfers in photosynthesis.

---

The absorption of solar energy and subsequent transduction to a chemical energy in the early stages of photosynthesis has a quantum efficency approaching unity.^1^ This efficiency and the ability of photosynthetic biomolecules to participate in electronic circuits^2^ makes them possible candidates for the development of photonic bio-hybrid nanotechnologies e.g., photo-biosensors^3-5^ and optoelectronics.^6, 7^ The natural photosynthetic membranes of green plants are found within chloroplasts and are termed “thylakoid membranes”. These membranes contain a large network of light-absorbing proteins, each protein containing a high density of pigments. Photon absorption by a pigment molecule (e.g., chlorophyll, carotenoid) produces an excited electronic state which can be transferred with high efficiency between the pigments within one protein and between nearby proteins. In plants, many copies of the protein “Light-Harvesting Complex II” (LHCII) form a large “antenna” system that performs rapid and highly efficient transfer of excitation energy to “Photosystem” (PS) proteins, which then perform photochemistry.^8-10^ Photochemistry in the PSII leads to a cycle of electron transport through the membrane, coupled to the unidirectional pumping of protons across the membrane, using several intermediate proton and electron carriers (see Figure 1A). This proton gradient is a temporary store of chemical and electrical energy, which can be harnessed to generate high-energy biomolecules (e.g., adenosine triphosphate and NADPH).^11^

**Figure 1.**
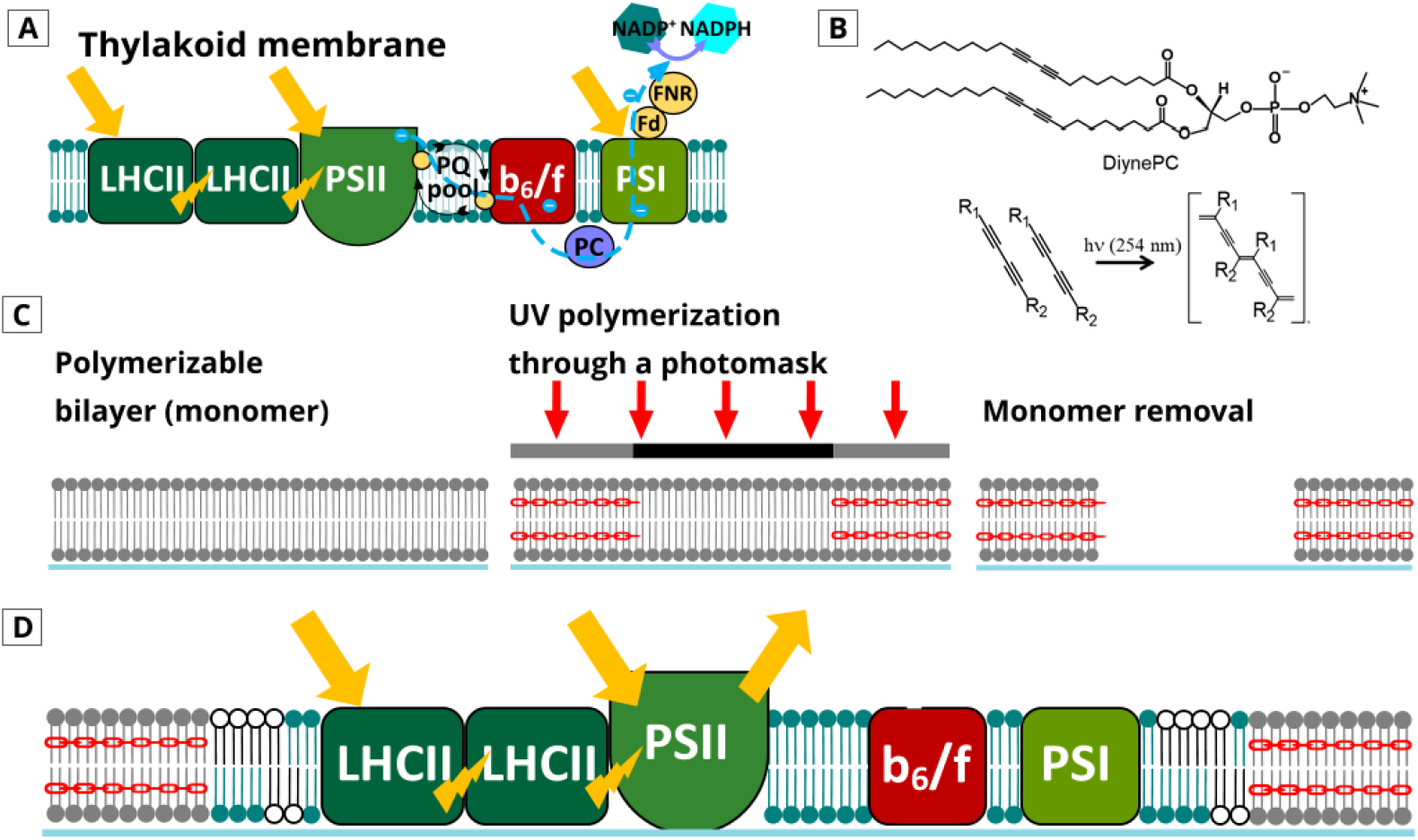
Concepts for designing model LH membranes, as reported recently.^14^ (A) Schematic of the natural thylakoid membranes and the energy transfer processes occurring. *Yellow arrows* represent absorption of light, *yellow bolts* represent inter-protein excitation energy transfer and *blue dashed lines* represent the electron transfer chain (simplified). (B) Chemical structure of the Diyne-PC and the photo-polymerization reaction. (C) Schematic of the how photo-polymerization is carried out through a photomask to generate array patterns, where only the regions of Diyne-PC exposed to UV become crosslinked (*red linkers* indicate polymerized lipids). (D) Schematic of the “hybrid membranes” within the polymer-lipid template.

The interaction between the photosynthetic machinery relies heavily on protein arrangement and the surrounding superstructure of the thylakoid lipid membrane.^12^ In the native system, LHCII and PSII are organized into “supercomplexes” which are located within stacked membranes, called “grana”.^13^ The overall stacked membrane arrangement provides a large surface area for incorporating many hundreds of pigments across >100 nm, creating a wide spatial and optical cross-section for the absorption of sunlight. The main antenna protein, LHCII, undergoes concentration-dependant quenching apparently based on the extent of LHCII-LHCII associations. Large-scale rearrangements of LHCII and PSII within grana can change the extent of protein-protein interactions, modulating the energy balance within the system and open/closing additional energy dissipation pathways.^15-21^ This effect is observed in spectroscopy of model systems (e.g., LHCII aggregates and proteoliposomes) and natural membranes (e.g., chloroplasts, plant leaves) as a descreased fluorescence intensity and shorter fluorescence lifetime, often termed “fluorescence quenching”.

Determining the concentration and relative positions of LH and PS proteins is therefore crucial for understanding both excitation energy migration and electron transport across membranes.^22-24^ Electron microscopy has been important for revealing the superstructure of photosynthetic membranes and the positions of proteins within membranes,^25-28^ however, it is time-consuming, expensive and cannot usually be performed under hydrated samples at room temperature. In contrast, Atomic Force Microscopy (AFM) allows the visualisation of LH proteins and PS complexes at relatively high resolution (∼1 nm laterally and ∼0.1 nm vertically) and can measure membrane samples under close-to-native conditions (ambient temperature, aqueous environment, etc.).^29-32^ Depositing thylakoid membranes which have been extracted from chloroplasts onto a flat, solid surface such as mica or glass can allow high-resolution AFM and spectroscopy to be performed and increase our understanding of the interactions within the native system. However, isolated and fragmented natural membranes are not necessarily an ideal test platform to test system functionality because of their heterogeneous composition and unstable nature.

The photophysical properties of LH proteins have also been studied utilizing nanoscale array patterns of proteins on solid surfaces^33-35^ and LH proteins incorporated into model membranes (proteoliposomes).^21, 36-41^ These model systems offer several advantages as platform to study the inherent physicochemical properties of the proteins, such as providing precise control over protein arrangement, known membrane composition and the incorporation of specific lipids to help maintain protein stability.^38, 42^ However, many of these models require extensive biochemical purification, chemical alteration of the protein or the support surface, and/or removal of the native lipids, any of which may affect the stability and photophysical state of LH and PS proteins.^37, 43-45^ Furthermore, these models are often limited to one or two types of protein which simplifies the complex interactions that are present in the native system, because of the procedural challenge of reconsituting multiple types of purified protein into a single artificial lipid membrane.

Therefore, there is a compelling need for a thylakoid membrane model that has an intermediate level of complexity: consisting of the full range of proteins found in the native thylakoid membrane, but with a greater control over membrane composition and amenability to high resolution microscopy (i.e., surface based). An ideal model system for the study of photosynthetic membranes would consist of a stable membrane on a solid support which contains the complete network of photosynthetic proteins embedded within a bilayer comprised of a native-like mixture of lipids. Supported Lipid Bilayers (SLBs) have regularly been used as a model system to understand the physical chemistry of lipids at the nansocale. SLBs can be “patterned” into an easily recognizable micro/nano array structure to allow more accurate analysis^46-48^, as well as providing a flexible morphology for the design of photo-electronic components. The reconstituion of LH and PS proteins into artificial lipid membranes can provide a model system to understand the photophysical and biochemical processes of photosynthesis and/or to inspire the design of new nanotechnologies.^49-51^ However, due to the high degree of curvature and the high protein density of native thylakoid membranes, it can be difficult to generate SLBs that are continuous across large areas.

Recently, the photo-polymerisation of diacetylene-phosphocholine (Diyne-PC) lipids into designer patterns by exposure to UV light through a photomask, has been used to form a highly stable array of SLB (see Figure 1B-1C).^52-54^ The empty Diyne-PC patterns have exposed lipid bilayer edges, which have a high free energy, promoting the formation of SLBs when the template is backfilled with a solution of lipid vesicles.^55^ The result is an array of discrete, high-quality SLBs which are corraled within a robust template.^52, 56^ Very recently, Morigaki et al. presented a new type of “hybrid membranes” by incorporating thylakoid components into SLBs within a Diyne-PC array-patterned template.^14^ This first characterization of the model used simple epifluorescence microscopy to visualize the incorporation of thylakoid proteins into the template patterns, at microscale resolution. The LH and PS proteins were able to diffuse laterally within the membrane, suggesting that the SLBs were high quality and functional assays suggested that the electron transport cycle was still somewhat active. However, this study was relatively qualitative and left many unknowns: (a) what are the dynamics of the hybrid membrane assembly process? (b) what is the structure of the lipid bilayers at the nanoscale? (c) what quantity of protein is incorporated into the membrane? (d) how efficient is excitation energy transfer between LH proteins (see Figure 1D)? In the current study, we attempt to answer all of these questions to test the efficacy of the hybrid membranes as a platform for studying photosynthetic protein interactions and developing new bio-hybrid light-harvesting materials.

## Results and Discussion

In the native system, proteins occupy ∼60-70% of the thylakoid membrane (by weight),^13^ leading to strong protein-protein associations that dictate the overall energy flow within the membrane. These tightly-packed LH proteins are known to have relatively quenched fluorescence compared to isolated LH proteins due to protein-protein interactions, therefore, changes to the protein arrangement will alter the degree of fluorescence quenching.^18, 21^ To assess these interactions between LH proteins we used fluorescent lifetime measurements which quantify the degree of quenching (from the excited state decay rate).^57^ Specifically, a laser-scanning fluorescence microscope was used to acquire images where each pixel has both fluorescence intensity and a time-resolved fluorescence spectrum, termed “Fluorescence Lifetime Imaging Microscopy” (FLIM).^18, 33^ This strategy was employed throughout the study, and allowed us the ability to observe the structural arrangement of our membrane samples correlated to their photophysical properties (at ∼300 nm resolution).

Firstly, a type of natural photosynthetic membranes was analyzed as a baseline to compare to our model system. Thylakoid membranes were extracted from plant chloroplasts following standard protocols and a basic spectroscopy characterization confirmed that these samples contained the expected proteins (LHCII, PSII, PSI, see Supplementary Figure S1). These “extracted thylakoids” were adhered to a hydrophilic glass coverslip to make them accessible to imaging via FLIM. Figure 2A shows a representative FLIM image obtained from the chlorophyll (Chl) fluorescence of extracted thylakoids, revealing many distinct objects which all appear to have similar, quite short fluorescence lifetimes of ∼0.5 ns. Note, all FLIM images have a color scale with fluorescence lifetime represented from blue (short lifetime) to red (long lifetime) and an intensity scale representing the total counts in each pixel. Control measurements show that there is minimal nonspecific fluorescence from any other sources or impurities (see Supplementary Figure S2), so we can be confident in the detection of chlorophyll fluorescence even where the signal is low. To analyze the Chl signal quantitatively, a fluorescence decay curve was generated by accumulating the photons collected from the whole image (Figure 2C, blue curve): this reveals a mean fluorescence lifetime <τ> = 0.40 ± 0.01 ns (N > 500 particles; ± standard deviation). This lifetime is in agreement with our ensemble spectroscopy data of the extracted thylakoids in solution (<τ> ∼ 0.5 ns) and is in good agreement with previous reports of LHCII and PSII within intact chloroplasts and leaves.^58-60^ This <τ> is much shorter than the lifetime known for isolated LH and PS proteins in detergent (∼ 4ns), as expected, due to the quenching effect of protein-protein interactions present in thylakoids. Overall, this shows that our FLIM data on LH membrane samples agree nicely with standard spectroscopy and that the deposition of the membranes on glass surfaces has no apparent impact on the sample.

**Figure 2.**
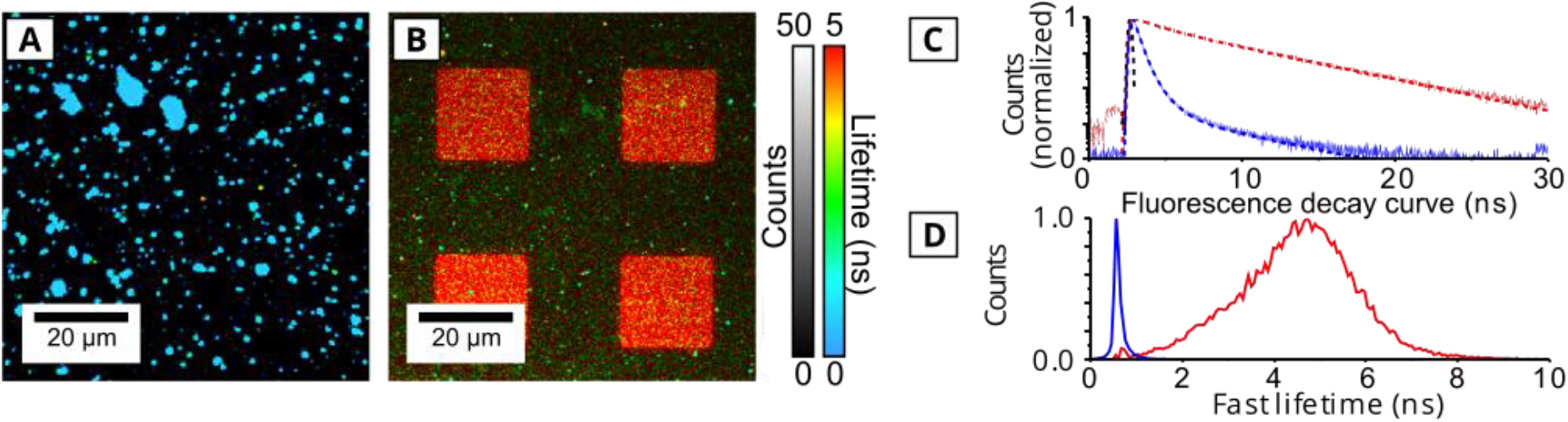
Analysis of extracted thylakoids versus hybrid membranes via FLIM. All samples were imaged in a buffer of 50 mM KH_2_PO_4,_ 10 mM NaCl, 20 mM MgCl_2_ and 330 mM sorbitol (pH 7.5). All images show selective excitation of chlorophyll at 485 nm and collection of fluorescence emission between 655 – 725 nm. (A) FLIM image of extracted thylakoid membranes adhered onto a hydrophilic glass surface (30 min incubation, then washed). (B) FLIM image of the “hybrid membranes” (30 min incubation of a polymerized Diyne-PC template with a 1:3 mixture of extracted thylakoid: DOPC liposomes, then washed). (C) Normalized fluorescence decay curves: showing raw data (*pale lines*) and fits (*dashed lines*) on log/linear axes (y/x). The *blue curve* is the ensemble of all counts from the intact thylakoids image in (A); the *red curve* includes the counts accumulated from 16 selected corrals (*box regions*) of hybrid membrane similar to (B). The mean amplitude-weighted lifetime, <τ>, was calculated by modelling the curve as a multi-exponential decay function (excellent fits were achieved for all data, with chi-squared values <1.1 and low residuals). (D) Frequency distribution histogram of the “FastFLIM” fluorescence lifetime, samples colored as in (C), normalized to a peak of 1 (“FastFLIM” is a high-throughput, analysis of the photon arrival time, see Methods).

Hybrid membranes were prepared in a two-stage process. In stage 1, templates of polymerized Diyne-PC on glass coverslips were generated by photolithography, as previously published^14^ and shown schematically in Figure 1C. The photolithogaphy generates a pattern based on the design of the photomask used; here, we chose a square-array grid pattern expected to produce an array of lipid bilayers with exposed edges providing large 20 x 20 μm corral regions of empty glass (see Supplementary Figure S3). In stage 2, natural and synthetic membranes were combined to fill the empty regions of the template and fuse with the exposed edges to form a corraled SLB. Specifically, Diyne-PC templates were incubated with an aqueous suspension of extracted thylakoids and synthetic lipid vesicles (DOPC), in a 1:3 weight/weight ratio, and the sample was washed with clean buffer solution. FLIM performed on these samples of “hybrid membranes” is shown in Figure 2B, revealing clear array patterns where the vast majority of Chl fluorescence is localized within the square corral regions defined by the template. These patterned hybrid membranes were highly reproducible, with similar dimensions and fluorescence intensity across multiple preparations (Supplementary Figure S4).

The Chl fluorescence lifetime measured here was much longer for these hybrid membranes compared to extracted thylakoids. The fluorescence decay curve generated from analyzing the photons accumulated from the box regions confirmed a slow decay process for hybrid membranes with <τ>= 4.06 ± 0.11 ns (*red curve* in Figure 2C). This value represents entirely “non-quenched” proteins (isolated LHCII in detergent has <τ> ∼4 ns^61^), in stark contrast to the short lifetime of extracted thylakoids. The long average lifetime suggests that the protein density in hybrid membranes must be sufficicently low that the protein-protein interactions found in the natural thylakoids (which reduce the fluorescence lifetime as discussed above) are relatively rare. Note, this long fluorescence lifetime was mainly observed inside of the corral region, with a minor sub-population of small particles with shorter (*bluer*) lifetimes observed on the surrounding framework. These short-lifetime particles are likely to be extracted thylakoids that have adhered to the top of the template and not merged with the synthetic lipid bilayers.

It is informative to assess the distribution of lifetimes within each sample, because this allows us to comment on the range of photophysical states within each membrane, rather than merely the average. To do this, a frequency distribution plot of fluorescence lifetime was generated for both samples by binning photons into appropriate time ranges, shown in Figure 2D. These distributions can be fit to Gaussian functions centred around 0.57 ns and 4.58 ns for the extracted thylakoid sample and the hybrid membrane sample, respectively. The width of the distribution was significantly narrower for extracted thylakoids than hybrid membranes (FWHM_extracted_ = 0.15 ns vs FWHM_hybrid_ = 2.31 ns). The broad distribution of lifetimes in the hybrid membranes suggests a variety of photophysical states of the LH and PS proteins, which could be caused by heterogeneity in the protein density or local density fluctuations. The dramatic increase in Chl fluorescence lifetime observed in both the fitted lifetime <τ> and frequency distributions, leads us to conclude that large-scale protein and lipid reorganisations occur during the hybrid membrane assembly. We hypothesise that the photosynthetic proteins become diluted significantly when thylakoid membranes merge with DOPC lipid bilayers.

To test this hypothesis, we attempted to quantify the change in protein density by careful analysis of the absolute magnitude of fluorescence emission between samples. The fluorescence intensity of hybrid membranes was compared to the fluorescence intensity of control samples containing a known amount of photosynthetic proteins, whilst taking into account changes in the level of quenching and keeping consistent acquisition parameters. A control sample of “LHCII proteoliposomes” was prepared from a defined quantity of purified LHCII and natural thylakoid lipids, as previously described^41^. The density of proteins in hybrid membranes was then calculated in terms of “LHCII equivalents” in a three stage calculation process. Firstly, the approximate number of counts per LHCII proteoliposome was calulcated based upon FLIM measurements (see Supplementary Figure S5 and Table S3). Secondly, the lipid/protein ratio was calculated from bulk spectroscopy and the estimated molecular packing (see Supplementary Table S4). Thirdly, the number of LHCII-equivalents per corral of hybrid membranes was calculated, based on FLIM data and the values from stages 1-2 (see Supplementary Table S5). Our best estimate for the protein content is 57,100 ± 5,500 LHCII-equivalents per corral (3,083,000 chlorophylls). Taking into account uncertainities, we estimate a possible range for the protein density of 84 – 451 LHCII/μm^2^ corresponding to 0.32 – 3.54 % of the total membrane area being occupied by photosynthetic proteins (best estimates of 143 LHCII/μm^2^ and 0.72%). Given that natural photosynthetic membranes are comprised of 60-70% protein by weight,^13^ these estimates are in agreement with our hypothesis that the hybrid membranes contain a relatively low concentration of proteins. In later sections, we assess if the apparent change in the photophysical state and reduced density affects the functionality and energy transfer within the system.

For our photosynthetic model, understanding the membrane formation process is useful for explaining its resulting composition and the photophysical state of the proteins incorporated. So, the kinetics of protein insertion into the hybrid membranes were studied in real-time using time-lapse FLIM. The intensity and lifetime of chlorophyll fluorescence during membrane assembly over sequential FLIM images are shown in Figure 3A (*brightness* represents intensity and *false-color* scale represents lifetime). Each image displayed represents the cumulative sum of all photons detected in a 20 s period, a minimal period which provides sufficient signal for analysis. Over the 30-minute duration of the experiment, the increasing Chl intensity observed is the combination of signal from two types of membranes, distinguished by their different fluorescence lifetimes: (i) inside the corrals the predominant signal was a relatively long fluorescent lifetime similar to that observed for the steady-state of the washed hybrid membranes (*red square* features in Figure 3A); (ii) across the image globular particles with a short fluorescent lifetimes became more numerous over time (large *blue/green* spots in Figure 3A), presumably representing extracted thylakoids that had not merged with the synthetic lipid bilayers. At later time points (after 300 s), the intensity due to extracted thylakoids continued to increase, ultimately obscuring the long-lifetime signal underneath. To quantify the rate of membrane deposition, we generated frequency distribution plots of photon fluorescence lifetimes for each 20 s timepoint. As anticipated from the FLIM images, we observed a bimodal distribution, consisting of a long lifetime peak and a short lifetime peak, consistent with the frequency distribution plots from the steady-state samples of hybrid membranes and extracted thylakoids (Figure 2D). The lifetime distribution for each timepoint was deconvoluted into two Gaussian populations, as shown in Figure 3C (acceptable fits were achieved for all timepoints, with R^2^ > 0.9). The peak amplitude of each Gaussian, representing the size of the sub-population, was plotted against time in order to determine the rate of deposition for each type of membrane, see Figure 3D. The amplitude for extracted thylakoids increased with time, suggesting a progressively increasing surface coverage (*blue line* in Figure 3D,). This signal may be expected to saturate after a sufficiently long time as the surface becomes completely covered by extracted thylakoids, but the deposition process was stopped before this point was reached. For hybrid membranes, the amplitude increased at a much faster rate reaching a maximum value at ∼500 s (*green line* in Figure 3D). It is likely that the saturation effect arises from the effect of filling the finite area within the corral regions. This is consistent with the model for a Langmuir isotherm,^62^ where the rate of material adsorption is proportional to the remaining free space on the substrate. In the early stages of the Langmuir model where there is a large amount of remaining free space the rate of deposition is almost linear, and then starts to slow down, and eventually saturate, as the surface becomes increasingly occupied.

**Figure 3.**
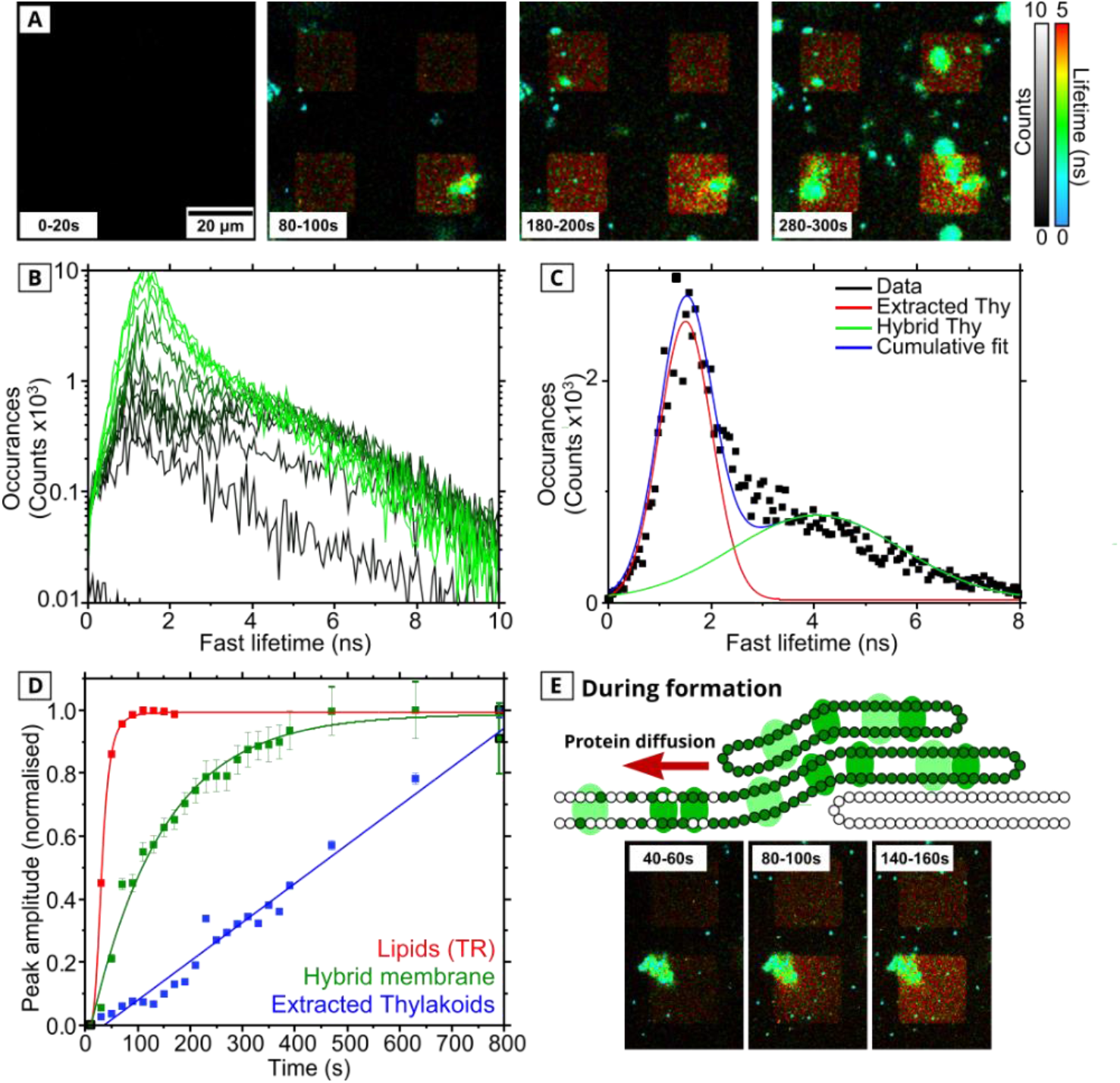
Analysis of the dynamic assembly processes occurring during hybrid membrane formation. (A) Time-lapse series of FLIM images during the formation of hybrid membranes. Each panel shows a 20-second acquisition during the real-time membrane assembly (using the same excitation and emission optics as in Figure 2). (B) Evolution of the frequency distribution of fluorescence lifetimes during the time series shown in (A). *Dark green* to *light green* colored curves represent the cumulative frequency distribution at increasing time-points of 20-40 s, 40-60 s, 60-80 s, 80-100 s, 100-120 s, 180-200 s, 220-240 s, 360-380s, 460-480s, 620-640 s, 780-800s, 940-960 s. (C) An example frequency distribution from a 20s snapshot (t = 220-240s), deconvoluted into two Gaussian functions (long-lifetime and short-lifetime). (D) Analysis of the growth of components of the hybrid membrane over time (normalized to its maximum value for display purposes). *Green*: photosynthetic proteins in the hybrid membrane, tracked through the long-lifetime peak amplitude from C. *Blue*: extracted thylakoids signal, tracked through the short-lifetime peak amplitude from C. *Red*: lipid accumulation, tracked through fluorescent lipids in a control sample (see Supplementary Figure S6). (E) Proposed schematic for the self-assembly of hybrid membranes with demonstrative fluorescence images: thylakoid membranes adhere to the top of a partially formed lipid bilayer and photosynthetic proteins migrate into the surface-adhered membrane via lipid bridges driven by concentration-dependent diffusion.

The introduction of synthetic lipids (DOPC) was essential for the formation of the hybrid membranes and thylakoids did not merge with the template in the absence of DOPC.^14^ To test the role of the synthetic lipids, a small amount of fluorescently-tagged lipids were incorporated into DOPC vesicles (0.5 % weight/weight Texas Red lipids), before mixing with extracted thylakoids. Time-resolved FLIM was performed tracking the fluorescence specific to the lipids during hybrid membrane formation and images were analyzed as above (*red line* in Figure 3C). By fitting each peak amplitiude curve to the Langmuir model (see Supplementary S8 for model derivation), the rate of lipid deposition was found to be 5.5 times greater than the rate of protein deposition (R_lipids_ = 0.039, R_proteins_ = 0.007). As a result, the lipid amplitude saturates much earlier (t_sat_ = 100 s for tagged-lipids, t_sat_ = 600 s for Chl-proteins) and FLIM images show the lipid fluorescence was homogenous across the corral region at t = 100 s, suggesting a close to, or completely, fluid DOPC bilayer at this time (see Supplementary Figure S6). Our results suggest that a bilayer of synthetic lipids is largely assembled inside the corral, before the majority of photosynthetic proteins have been incorporated into the membrane: in fact, the amplitude representing proteins assembling in hybrid membranes increases another two- to three-fold after the lipid signal has saturated (compare *green* vs *red* curve at t=100 s in Figure 3D). In examples where particularly large extracted thylakoids had adhered on top of a corral, chlorophyll fluorescence appeared to spread from these particles into the surrounding area, see images in Figure 3E. We hypothesize that extracted thylakoids (*blue-green* particles) adhere on top of the nascent DOPC bilayer and act as resevoirs, from which photosynthetic proteins undergo concentration-driven diffusion into the spreading hybrid membrane, as proposed in the cartoon in Figure 3E. It is unlikely that proteins could transfer vertically between stacked lipid multilayers (because this would expose hydrophobic portions of the protein to the polar solvent which would be thermodynamically unfavourable), so it seems probable that proteins and thylakoid lipids migrate laterally between lipid bridges that connect the ruptured thylakoids to patches of putative hybrid membrane. This interpretation is in agreement with other studies which observed lipid and cofactor diffusion between multilayers of stacked model membranes^63^ and bears similarities to the dynamic protein rearrangements which occur in natural thyalkoids.^12, 24^

After hybrid membrane formation and stabilization (washing to remove any loosely associated material), photobleaching experiments suggested that the majority of proteins and lipids were able to diffuse laterally within the membrane (Supplementary Figure S7), suggesting that the lateral motion of proteins/lipids was not significantly impeded by interactions with the substrate. The high fraction of mobile proteins also suggest that the method of hybrid membrane formation could selectively sort proteins in one-orientation. For example, PSII is asymmetric in its protrusions from the lipid bilayer (∼4 nm on the lumenal side, compared to ∼1.7 nm on the stromal side),^30^ and may be immobile when the bulky (lumenal) side experiences friction with the surface or may be sterically excluded from scenarios where the lumenal side is appressed with the substrate. Another possibility is that only mobile proteins are able to diffuse along lipid bridges from thylakoids into hybrid membranes that are supported by the substrate.

Next, the structural continuity of the membrane was assessed to test for any imperfections which could affect the function of the hybrid membranes as a model. An instrument combining AFM with FLIM was used to record nanoscale topography maps spatially correlated to multi-channel fluorescence data. Spectral and temporal selectivity of the FLIM allowed us to define two separate FLIM channels: (i) the “Chlorophyll (Chl) channel” defined as the combination of selective Chl excitation and a detector optimized for Chl detection, (ii) the “Diyne-PC Channel” optimized for the excitation and emission of the intrinsic fluorescence of the polymerized lipid template (note, the spectral overlap between the two fluorescence channels was negligible, see Supplementary Figure S2 and Table S1). These two fluorescence channels were probed simultaneously using a pulse-interleaved excitation mode, with AFM topographs acquired on the same region immediately after, images shown in Figure 4A-B. The AFM height profile in Figure 4C (*red line*) revealed a 4.81 ± 0.07 nm height from the polymerised lipids to the base of the empty corral, in excellent allignment with the fluorescence intensity profile which drops from ∼75 counts to ∼0 counts over the same region (Figure 4C, *blue line*). For the “empty” lipid template, the background signal in the chlorophyll FLIM channel was approximately zero across the entire image, as expected (Figure 4C, *green line*). After the formation of the hybrid membranes, there was largely homogenous Chl fluorescence within the square corral regions with no resolvable defects at this scale. The increase in the Chl fluorescence intensity (Figure 4D, *green line*) corresponded with a step change in the AFM height to a mere 0.19 ± 0.08 nm (Figure 4D, *red line*). Thus, the thickness of the thylakoid membrane was inferred to be 4.62 ± 0.15 nm.

**Figure 4:**
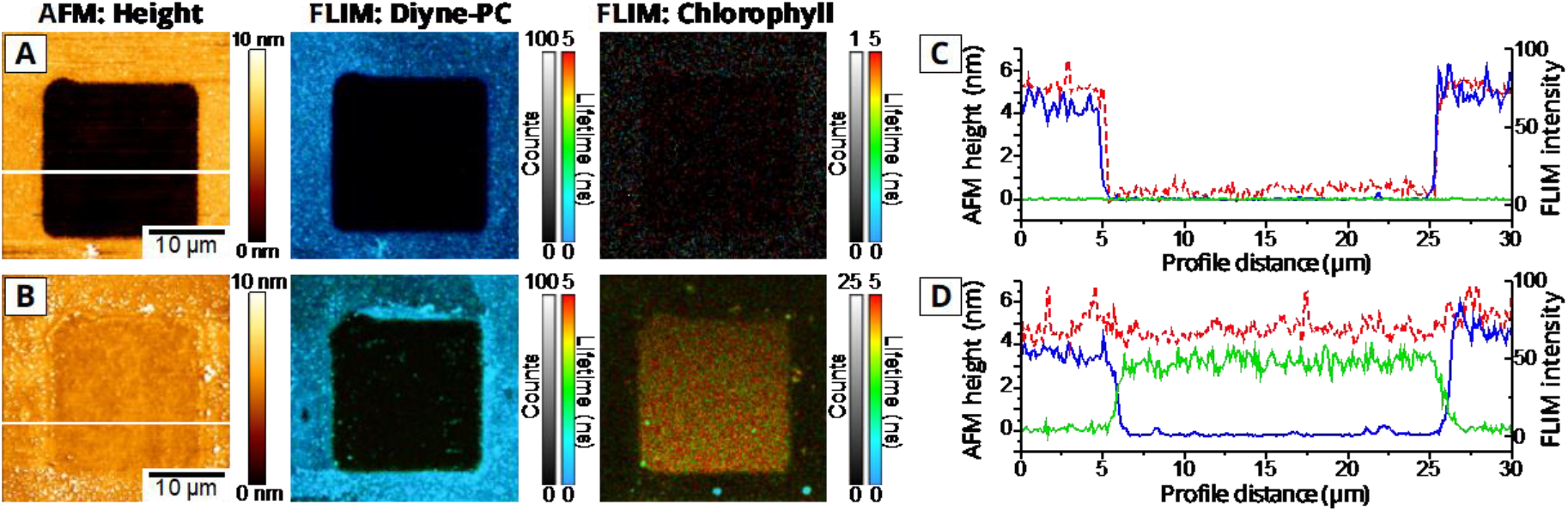
Analysis of hybrid membranes by correlated FLIM and AFM measurements. For (A) and (B) the *left panel* is an AFM topograph, the *center panel* is the “Diyne-PC FLIM channel” (i.e., optimized to detect the polymerized lipid by using excitation at 485 nm and collection of emission between 505-535 nm), the *right panel* is the “Chl FLIM channel” (i.e., optimized to detect the chlorophyll fluorescence from LH and PS proteins by using excitation at 640 nm and collection of emission between 672-696nm). (A) Correlated FLIM+AFM data showing a single square of the polymerized lipid “empty” template. The minimal signal in the Chl FLIM channel is statistically indistinguishable from detector noise. (B) Correlated FLIM+AFM data showing a similar region as in (A), but after it was “backfilled” with the extracted thylakoids and DOPC liposomes to form the hybrid membrane within the corrals. (C) and (D) show profiles drawn across the region indicated with a *white line* in (A) or (B), respectively: showing the AFM height (*red, dashed*), FLIM intensity from Diyne-PC (*blue*) and FLIM intensity from Chl (*green*). The Chl intensity is displayed after multiplication by a factor of 3, for comparison purposes. Higher magnification FLIM+AFM data is shown in the Supplementary Figure S8.

To reveal the smaller features which are expected, such as nanoscale protein protrusions and nanoscale membrane defects which may arise from deposition processes, a higher magnification structural mapping was performed, using a standalone AFM instrument which has higher spatial resolution. Measurements on a control sample formed entirely from DOPC lipids show a contiguous and defect-free membrane within the polymerized template (see Supplementary Figure S9), confirming the quality of our synthetic lipid vesicles. AFM measurements of hybrid membranes reveal a more complex topography, which we attribute to incorporation of thylakoid proteins and lipids. The hybrid membrane thickness was confirmed as ∼4.5 nm by direct comparison of the same corral before and after backfilling with hybrid membrane (Figure 5A vs 5B and *inset graph*). The high resolution AFM data also revealed that ∼10% of the membrane is occupied with multiple small pores (Figure 5D vs Figure 5C), with a lateral scale of ∼100 nm and 4.45 ± 0.62 nm in depth (n= 10). The depth of these pores is in agreement with our inferred membrane thickness and concurs with published values for a DOPC bilayer (∼4.5 nm).^64^ At the base of many of these pores, static particles were observed, ranging from 4 to 10 nm in height. These particles could be classified into subpopulations that roughly agree with the dimensions expected for LH and PS proteins^30^ (*P1*-*P3* in Figure 5F) and may represent a variety of photosynthetic proteins which have become immobilized at these pores (further statistical analysis of these populations is shown in Supplementary Figure S10). One possibility is that these pores form when thylakoids which are loosely associated and have partially fused with the SLB are stripped away from the surface, pulling away sections of the membrane and leaving some LH/PS proteins stuck to the surface (shown schematically in the cartoon in Figure 5G). To improve our hybrid membranes in future work, it may be possible to “heal” such defects in lipid bilayers^65^ to provide increased membrane continuity, by a secondary addition of lipid vesicles which may spread into the pores and merge with existing lipid bilayers.^48^

**Figure 5.**
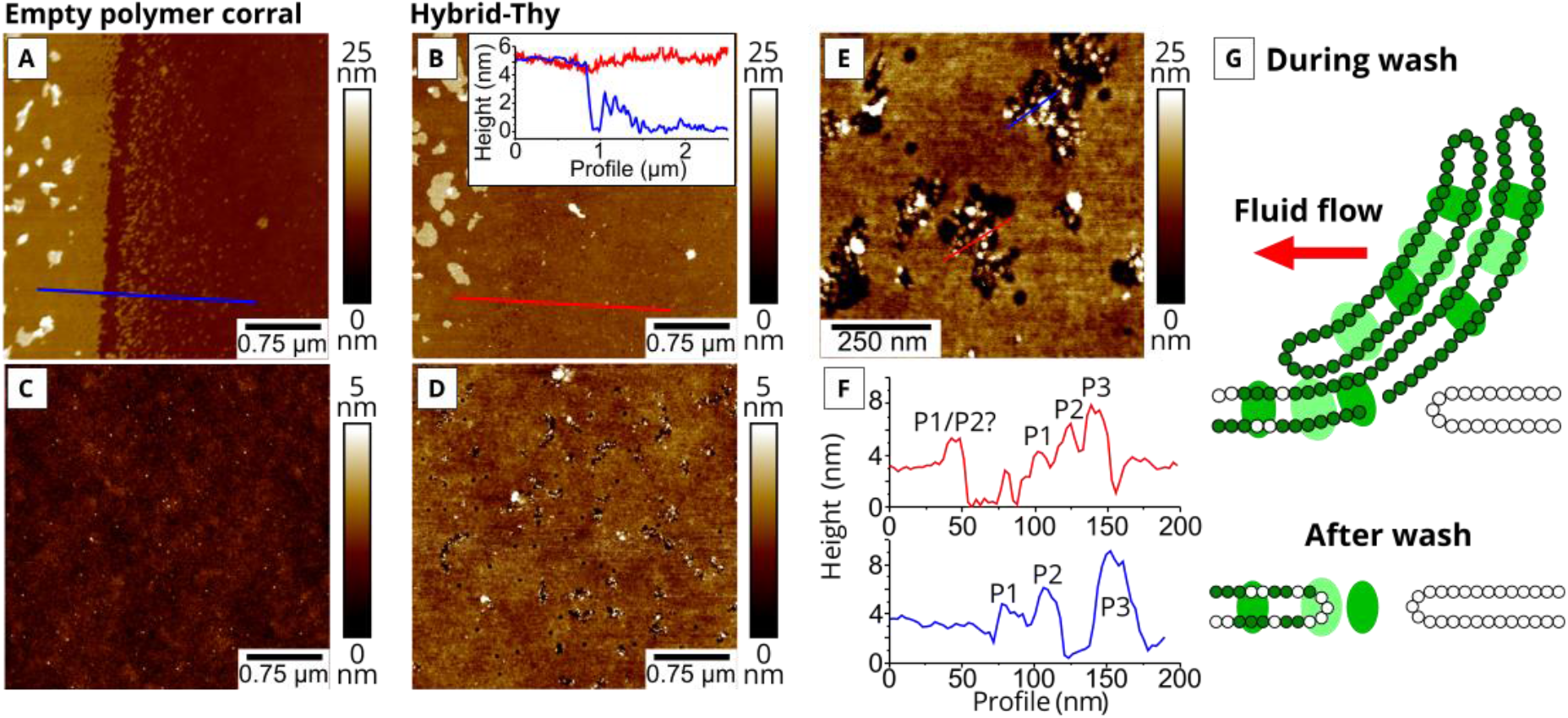
High resolution AFM analysis of empty templates versus hybrid membranes. (A) AFM of the step from an empty template to the base of the corral. (B) The edge of the template from (A), after the corral had been “backfilled” with the hybrid membrane (thylakoids and DOPC lipids). The *inset graph* is a height profile across this step, plotted for the backfilled corral (*red*) against the empty corral from (A) (*blue*). (C) An area inside of the empty corral at medium magnification, showing the glass substrate with its relatively smooth surface. RMS roughness = 0.353 nm (analysis of a 10 x 10 μm region). (D) An area inside the corral after backfilling, showing the structure of the hybrid membrane and membrane pores that become visible at this resolution. (E) An area of the hybrid membrane at high magnification where particles can be observed within the pores. (F) Height profiles from the *red* and *blue lines* shown in panel (E), showing three possible types of proteins within the bilayer (denoted *P1, P2* and *P3*). (G) A schematic for a possible mechanism for the formation of membrane pores. See Supplementary Figure S11 for a full gallery of AFM topographs.

In addition to the particles observed within pores, higher magnification AFM topographs (Figure 5E), shows many particles embedded within the lipid bilayer. To identify whether these protrusions are likely to be photosynthetic proteins, the density of particles was calculated for hybrid membranes in comparison to control samples containing no proteins (as a baseline estimate for the number of particles that may arise from subtrate imperfections or contaminants). A careful manual analysis of particle density in hybrid membranes showed significantly higher numbers (77.0 particles/μm^2^), than control samples of bare glass surface (13.8 particles/μm^2^) and the surface of DOPC lipid bilayer (9.7 particles/μm^2^). This suggests that >80% of the particles observed in hybrid membranes by AFM are photosynthetic proteins, leading to an estimated protein density of ∼60 proteins/μm^2^. This approximated protein density from AFM measurements was on the same order, but lower, than our previous estimates of protein density calculated via fluorescence intensity (estimations of ∼60 proteins/μm^2^ from AFM compared to 84-451 proteins/μm^2^ from fluoresence). This difference can be explained by considering that only static proteins can be observed by AFM, and that any highly mobile molecules would be “invisible” to the slow raster speed of the AFM probe. Therefore, the protein density estimated by AFM may only represent a small minority of the total population (∼10-15%), in agreement with the significant degree of lateral diffusion of proteins observed with photobleaching measurements (Supplementary Figure S7).

Finally, we attempted to assess the photosynthetic activity of the hybrid membranes, specifically, the transduction of excitation energy into electron transport (photochemistry). If this functionality is even partly retained in hybrid membranes, then this model system could be used to investigate these fundamental processes, or, due to the ability of PSII to donate electrons to downstream inorganic systems, may have applications in future photo-electronic technologies.^66, 67^ Multiple studies have proposed that, by selectively switching on or off portions of the electron transfer energy process, various photochemical inhibitors can give an indirect measure of the activity of PSII.^68^ Therefore, we performed a “photochemical assay” on hybrid membranes by monitoring changes to the chlorophyll fluorescence intensity (or lifetime), in response to these stimuli. In hybrid membranes as prepared, the water-soluble proteins responsible for electron transport from PSII to other proteins are likely to be missing (see Figure 6A). In this scenario, absorbed energy from PSII would be primarily released as Chl fluorescence. In the first stage of the assay to test the system, an exogenous electron acceptor, DMBQ, is introduced to the membranes at a relatively high concentration, to replace the natural electron carriers (PQ) which are likely to be saturated.^69-71^ If DMBQ successfully accepts electrons from PSII, the level of Chl fluorescence should be reduced in its presence compared to its absence, because excitation energy can be used to eject electrons rather than being re-emitted. In the final stage of the assay, “hydroxylamine” can be added as an aqueous solution and is known to increase the Chl fluorescence again (see Figure 6A).^72-74^ Hydroxylamine is a small highly reactive compound, reported to affect various cofactors within PSII, disrupting the oxygen-evolving complex and inhibiting the electron transfer cycle.^75^ The photochemical assay described above was performed on hybrid membranes and control samples and characterized by FLIM. Figure 6B(i)-(ii) show that the Chl fluorescence intensity is indeed significantly quenched after the addition of DMBQ, to 55 % (± 13 %) of its original intensity. The additional decay mechanism resulted in the decreased fluorescence lifetime within the corrals of hybrid membrane (from *red* to *green* on the false colour FLIM scale). Upon the addition of hydroxylamine, the fluorescence intensity recovered back to 97 % (± 17 %) of its initial intensity, see Figure 6B(iii). The trends for fluorescence intensity and for fluorescence lifetime of hybrid membranes are shown as *green lines* in the graphs of Figure 6C and 6D, respectively.

**Figure 6:**
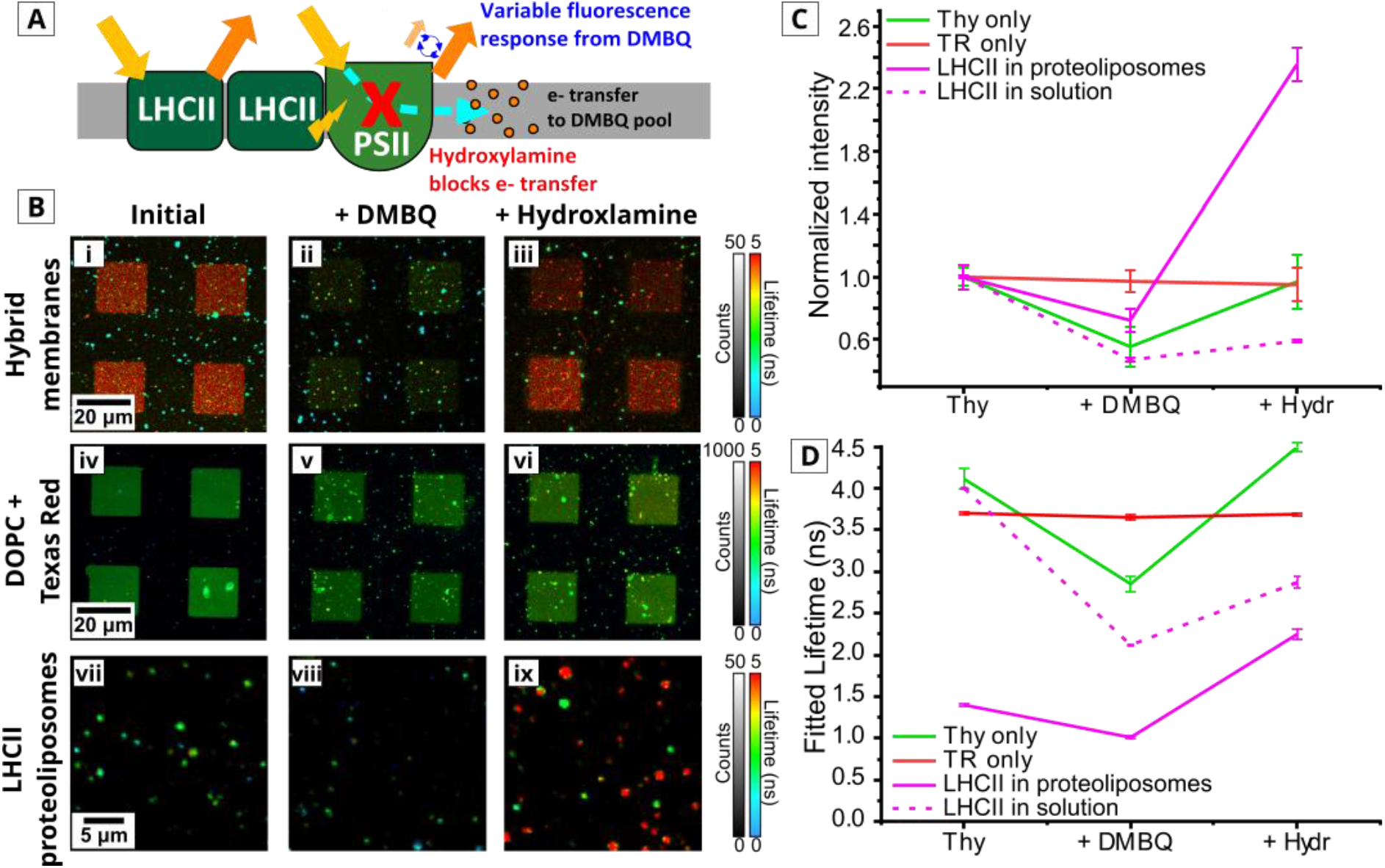
Quantification of possible photochemical activity using hybrid membranes and FLIM. (A) A schematic showing the possibilities for photon absorption, energy transfer and electron transfer in a hybrid membrane (various biomolecules after PSII not shown for simplicity). After DMBQ addition, excitation energy will drive electron transport from PSII to DMBQ, instead. After the addition of hydroxylamine, the PSII is no longer able to perform photochemistry and electron transfer is blocked, resulting in increased Chl fluorescence to dissipate the excess excitation energy. (B) FLIM measurements of hybrid membranes, and DOPC+TR lipid membranes and LHCII proteoliposomes, before and after the addition of 0.5 mM DMBQ and, after the addition of 0.5 mM hydroxylamine, as labelled. The FLIM instrument was set up with 640 nm excitation and 672-696 nm emission for (i)-(iii) and (vii)-(ix), or with 561 nm excitation and 590-650 nm emission for (iv)-(vi). Spectral overlap between these FLIM channels was minimal (see Supplementary Table S2). (C) Graph showing the normalized fluorescence counts of each sample plotted against the different experimental conditions from (B) (averaged over 4 fields of view). (D) Graph showing the mean fluorescence lifetimes for each sample against the experimental conditions from (B) (averaged over 4 fields of view). The raw data for the cuvette-based spectroscopy of LHCII proteoliposomes and LHCII in detergent solution is shown in Supplementary Figure S12.

The fluorescence lifetime was reduced from 4.11 ± 0.12 ns initially to 2.85 ± 0.09 ns upon the addition of DMBQ, before recovering to 4.49 ± 0.05 ns upon the addition of the hydroxylamine. This final lifetime is longer than in the initial system (4.11 ± 0.12 ns) and could be due to changes in the configuration of pigments within PSII after the addition of hydroxylamine.

A series of control samples were studied to assess the specificity of this assay on photosynthetic proteins and for probing electron transfer. DOPC lipid membranes containing the fluorophore Texas Red (without any proteins) showed typical images of patterned membranes (see Figure 6B(iv)-(vi)), with no significant change at any stage of the photochemical assay of either the fluorescence lifetime or the fluorescence emission intensity. This showed that DMBQ does not cause quenching of this chromophore and hydroxylamine does not affect it either (*red lines* in Figure 6C-D). Control samples of LHCII proteoliposomes deposited onto glass and LHCII in detergent (representing the quenched or non-quenched states of LHCII, respectively) were assessed as samples which contained Chl but not PSII (solid and dashed *magenta lines* in Figure 6C-D and Supplementary Figure S12). Therefore, these samples are examples of photosynthetic proteins that lack the electron transfer functionality inherent in PSII that this assay is expected to probe. Surprisingly, the DMBQ caused quenching of the fluorescence intensity and lifetime in both LHCII-only samples, followed by the subsequent de-quenching by hydroxylamine. LHCII in proteoliposomes showed significantly less DMBQ-induced quenching and more hydroxylamine-induced de-quenching compared to LHCII in solution, presumably because LHCII in proteoliposomes started in an already heavily quenched state. These positive results for LHCII suggest that the DMBQ/hydroxylamine assay is not specific for detecting electron transfer, as LHCII does not normally release electrons. A direct “collisional quenching” mechanism where DMBQ interacts with Chl excited states, either diffusing into the membrane or through the aqueous solution, to cause de-excitation seems possible.^76^ The finding that DMBQ did not affect the Texas Red (tagged to lipids) may be due to the different excited state energies of this fluorophore as compared to Chl. Hydroxylamine, also, must not be specific for binding to and disrupting PSII suggesting that this chemical could have the potential to react with and disrupt any Chl-containing protein. The increased Chl fluorescence lifetime of >4 ns suggests that hydroxylamine may either cause some sort of disaggregation of Chls which increases the intra-protein Chl-Chl distance, or a chemical change to the Chls which decreases the dipole-dipole coupling.

These photochemical experiments demonstrate one major advantage of utilizing a range of model membrane systems which are assembled from the desired components: namely that we can perform measurements on membranes containing only one type of protein to illuminate potential issues. Our comparisons of LHCII in a quenched and non-quenched state, also demonstrate that the initial photophysical state of the proteins can significantly alter the outcomes/interpretation of functional assays. A virtue of the use of FLIM (over intensity-only measurements) is that we can quantify such photophysical states more definitively to give confidence in our interpretations. Despite the apparent lack of specificity of the photochemical assays, they show that our hybrid membranes contain “active” Chl which responds to chemical modifications in a very different manner to other fluorophores, such as Texas Red.

## Conclusions

Our hybrid membranes have distinct advantages when compared to model membranes that have previously been developed. Alternative models which are formed by “bottom-up” approaches generally use purified proteins and lipids to exert control over composition, but this comes at the expense of missing many important components and a resulting simplification of the complex interactions observed in natural membranes.^21, 36-40, 42, 49-51^ Our hybrid membranes offer an intermediate situation, where natural membranes act as starting material so that all of the natural LH and PS proteins are potentially available for analysis, albeit at a lower density (approx. 1% of the membrane area). This model system maintains a native-like lipid environment and allows high rates of lateral diffusion of the proteins. The confined regions of hybrid membrane with their stable structure on a solid support surface and clear array pattern, together, provide a system that is amenable to high-resolution microscopy (much larger and flatter than natural thylakoids). There are many possible avenues which could be pursued in future studies to increase the protein concentration of the hybrid membranes to better represent the native system, such as altering the concentrations of the starting material,^14^ or by using techniques established in the SLB community to direct the diffusion of membrane proteins.^77, 78^

The excellent amenability of hybrid membranes to FLIM and AFM allowed us to explain their resulting structure and photophysical properties by imaging the self-assembly of lipids and photosynthetic proteins onto the solid surface in real time. In future, it may be possible to take advantage of the self-assembly mechanism to introduce additional lipophilic components, in order to enhance the photosynthetic activity to generate idealized designs of membranes (e.g., broadening the spectral range with additional pigments^41^). We found that the hydrophobic edge of the lipid corral and the admixture of DOPC liposomes promoted the formation of flat and contiguous membranes, even with the challenging starting material of extracted thylakoids (possibly due to an energy minimization considerations^56, 79^). This suggests that the polymerized lipid template could be used to support the formation of SLBs from a range of biological membranes that are otherwise difficult to study (high-curvature, protein-dense). A key advantage of using model membrane systems is that specific proteins of interest can be investigated, as shown in our application of our hybrid membranes (and proteoliposomes) to photochemical assays. This revealed new challenges in accurately determining electron transport using DMBQ and hydroxylamine, suggesting that the assays used in the photosynthesis community may have to be reassessed.^72-75^

## Materials and methods

### Preparation of extracted thylakoid membranes, purified LHCII, and lipid/protein vesicles

Thylakoid membranes were isolated from spinach (*Spinacia oleracea*) as described by Morigaki and co-workers.^14^ Briefly, this involved macerating leaves at 4°C, disruption of the chloroplasts by passing them through a high-pressure vessel and recovery of thylakoid membranes in an aqueous buffer (50 mM KH_2_PO_4_, 10 mM NaCl, 2 mM MgCl_2_, 330 mM sorbitol, pH 7.5). Absorption specstropy confirmed that the membranes contained the expected optically-active proteins (LHCII, PSII, PSI, see Supplementary Figure S1). These “extracted thylakoids” were used to form hyrbid membranes within a few days or were flash-frozen with liquid nitrogen and stored at −80°C. Purified trimeric LHCII (required for control measurements) was extracted and purified from spinach leaves following established procedures using detergent (*n*-dodecyl-*alpha*-*D*-maltopyranoside), sucrose density gradient sedimentation and size-exclusion chromatography (purity was confirmed by denaturing and native gel electrophoresis).^18^ LHCII proteoliposomes were prepared by combining specified quantifies of lipids, detergent and purified LHCII protein in an aquesous buffer and then inducing the self-assembly of membranes by removing the detergent using Biobeads, as described previously.^41^ Lipid vesicles, required for hybrid membranes formations, were formed from high purity 1,2-dioleoyl-*sn*-glycero-3-phosphocholine (DOPC) lipids following standard probe sonication procedures, with Texas Red lipids (Texas Red 1,2-dihexadecanoyl-*sn*-glycero-3-phosphoethanolamine) included only when tracking lipids was required.

### Preparation of polymerized lipid templates and hybrid membranes

The polymerized lipid templates were prepared as described in several previous publications.^52, 53^ Briefly, lipid bilayers of 1,2-bis(10,12-tricosadiynoyl)-*sn*-glycero-3-phosphocholine (Diyne-PC) were deposited onto substrates by vesicle spreading and then polymerization was conducted by UV irradiation using a mercury lamp, using very careful control over power delivered, process temperature and presence of oxygen. Substrates patterned with polymerized Diyne-PC could be stored in water for weeks at room temperature. Immediately before use, patterned substrates were dried with nitrogen and placed into a microscopy sample holder as desired (either, ultrathin adhesive imaging spacers to confine the sample droplet or the AFM OEM coverslip holders). Extracted thylakoids and DOPC vesicle suspensions were combined in a 1:3 weight/weight ratio and added to the substrate at a final concentration of 0.68 mM DOPC. After 30 min incubation, samples were rinsed with copious buffer solution and were ready for microscopy.

### Atomic Force Microscopy (AFM)

Standalone AFM was performed under aqueous buffers using a Bruker Dimension FastScan and PEAKFORCE-HIRS-SSB probes (Bruker AFM Probes) in PeakForce tapping mode. Parameters were optimised whilst imaging to minimise applied forces of <0.2 nN, typically scanning at 2-4 Hz and 1024 x 1024 pixels. Topographs were processed and analysed using Nanoscope Analysis Software (v1.9). For combined FLIM+AFM, the AFM imaging used a JPK NanoWizard 4 driven by a Vortis Advanced control station. A JPK Tip Assisted Optics stage was used for a sample-scanning configuration, so that once the FLIM laser spot and AFM probe were aligned they remained in a fixed position to ensure consistent correlation between the two systems and minimal noise.

### Fluorescence microscopy

Fluorescence Lifetime Imaging Microscopy (FLIM) was performed using a Microtime 200 time-resolved fluorescence microscope (PicoQuant GmbG). This system used an Olympus IX73 inverted optical microscope as a sample holder with light passing into and exiting various filter units for laser scanning, emission detection and timing electronics. Excitation lasers (picosecond pulsed sources) were driven in Pulsed Interleaved Excitation mode by a PDL 828 Sepia II burst generator module. The pulse width for the LDH 485 nm, LDH 561 nm, and LDH 640 nm lasers, were 90 ps, 70 ps and 90 ps respectively. Detector 1 was a Single-Photon Avalanche Diode and detector 2 was a hybrid Photomultiplier Tube. Specific dichroic mirrors and emission filters, as described in the text, were used to define the emission channel wavelength range. An excitation fluence of 0.012 mJ/cm^2^ was used which allowed sufficient fluorescence signal whilst limiting any singlet-singlet annihilation events (optimization shown in Supplementary Figure S13 and Tables S6 and S7). Similarly, we ruled out any significant effects due to the presence or absence of dissolved oxygen in the buffer on the photophysical properties the proteins (see Supplementary Figure S14). Analysis of all FLIM data was performed with SymPhoTime software (PicoQuant).

## Supporting information

Graphic abstract

## Acknowledgments

This collaboration between Leeds and Kobe was supported by an International Exchanges Cost Share award from The Royal Society UK (IEC\R3\183029) and Japan-UK Research Cooperative Program award from the Japan Society for the Promotion of Science (JPJSBP120195707). S.A.M. was supported by a Biotechnology and Biological Sciences Research Council (BBSRC, UK) studentship, award number BB/M011151/1. A.M.H. was supported by an Engineering and Physical Sciences Research Council (EPSRC, UK) studentship, award number 1807029, and an EPSRC program grant, EP/ J017566/1 “CAPITALs”. S.D.C. was supported by an EPSRC grant, EPSRC EP/J017566/1. S.D.E. was supported by funding from the Medical Research Council (MR/M009084/1) and the EPSRC (EP/P023266/1). T.Y. and K.M. were supported by the Japan-UK Research Cooperative Program (JPJSBP120195707) and a Grant-in-Aid for Scientific research (#19H04725) from Japan Society for the Promotion of Science (JSPS). T.Y. and K.M. thank Dr. Daisuke Takagi (Setsunan University, Japan) for discussion and support in purification and assays of thylakoid. P.G.A. was supported by a University Academic Fellowship (University of Leeds). The PicoQuant FLIM instrument at Leeds was acquired with funding from a BBSRC award number BB/R000174/1.

## Notes

### Competing Interest Statement

The authors have declared no competing interest.

