## Supporting Information for "Understanding the photophysics and structural organization of photosynthetic proteins using model lipid membranes assembled from natural plant thylakoids"

#### 1. Ensemble absorption and fluorescence spectroscopy of extracted thylakoids show they contain all the expected photosynthetic components

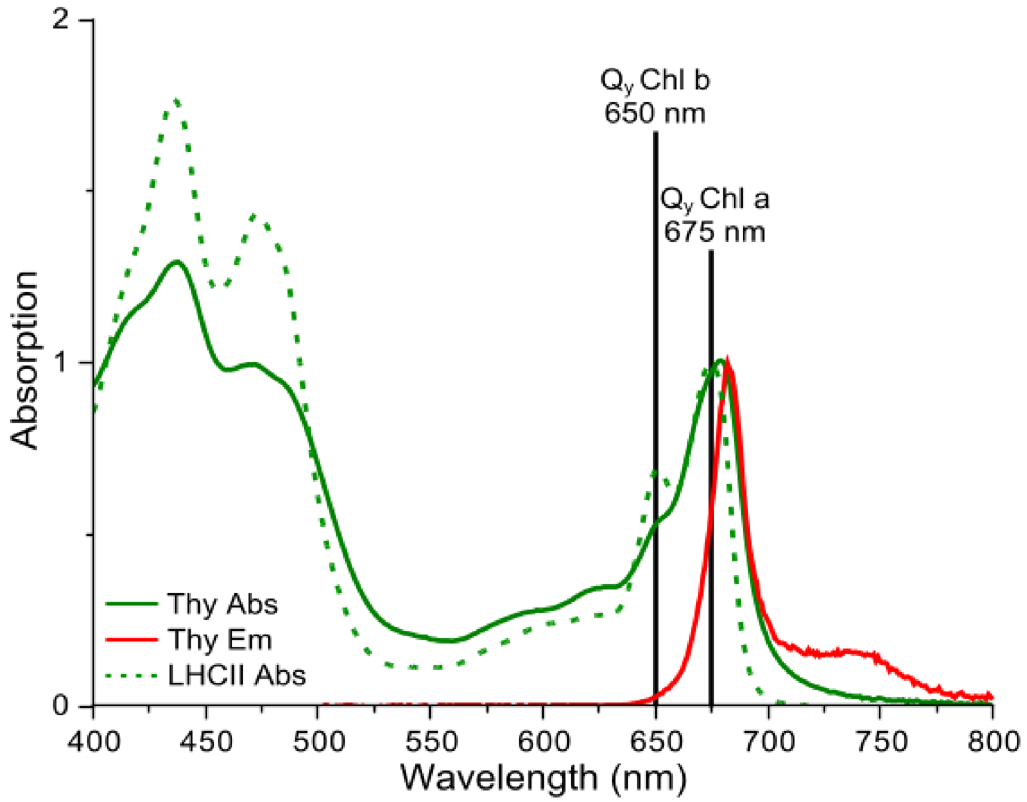


**Figure S1**

Emission (*solid red*) and absorption (*solid green*) data for extracted thylakoids and absorption data for LHCII (*dashed green*). Cuvette-based absorption spectroscopy was performed using an Agilent Technologies Cary 5000 UV-Vis-NIR absorption spectrophotometer. Cuvette-based fluorescence spectroscopy was performed using an Edinburgh Instruments FLS980 system, using excitation at 473 nm and collecting emission from 500-800 nm (2 nm and 1 nm bandwidth excitation and emission slits, respectively). The absorption spectra from both samples share some features, with similar peaks but at subtly different maximum wavelengths and with varying peak intensities. As expected, the overlapping peaks due to carotenoids and the chlorophyll Soret bands are found between 400-500 nm and the known $Q_y$ bands for chlorophylls at 650-700 nm.

Taking a closer look at the LHCII absorption spectrum (*green dashed line*), there is a clear peak at precisely 675 nm and a lower intensity peak at precisely 650 nm, representing Chl *a* and Chl *b* $Q_y$ transitions, respectively. The ratio of peak heights between Chl *a* and Chl *b* is as expected for LHCII and the subtle shoulder at 475 nm is indicative of the trimeric form of LHCII. These peak wavelengths and intensities are in excellent agreement with previous reports of the optical properties of spinach LHCII.[1-4] Gel electrophoresis of the purified protein shows the expected bands, as previously reported by our group.[1, 2]

In comparison, the extracted thylakoids absorption spectrum (*solid green*) is more complex, with contributions from LHCII, PSII and PSI. There is a peak centred at ~680 nm which is significantly broader than for LHCII and there is only a shoulder at ~650 nm, rather than a distinct peak. In summary, this spectrum is very similar to published reports for extracted thylakoids.[5] In detail, the well-established features of plant thylakoids, are explained by the following: (i) PSI and its antenna has maximal Chl *a* absorption at ~682 nm as compared to the PSII at a maximum of ~677 nm, explaining why this peak is found at longer wavelength compared to isolated LHCII alone, (ii) the PSI peak is known to extend further into the red due to its more numerous low-energy chlorophylls, explaining the observed broadening of the peak towards the red end of the spectrum, (iii) furthermore, PSII has much lesser Chl *b* than LHCII, and PSI has even less again, explaining the reduced peak at ~650 nm. This is in good agreement with reports by Caffarri and co-workers[5] and others.

The extracted thylakoids fluorescence emission spectrum (*solid red*) has one major peak at ~682 nm and broad "red tail" between 700-750 nm represents emission from low energy Chl *a* (and its extended vibrionic manifold)[6, 7] indicating that the highly connected chlorophyll network across photosynthetic proteins has been maintained in the preparation of the "extracted thylakoids" sample, again, as we would have expected.

#### 2. Quantification of spectral overlap between targeted FLIM channels (confirming minimal nonspecific fluorescence)

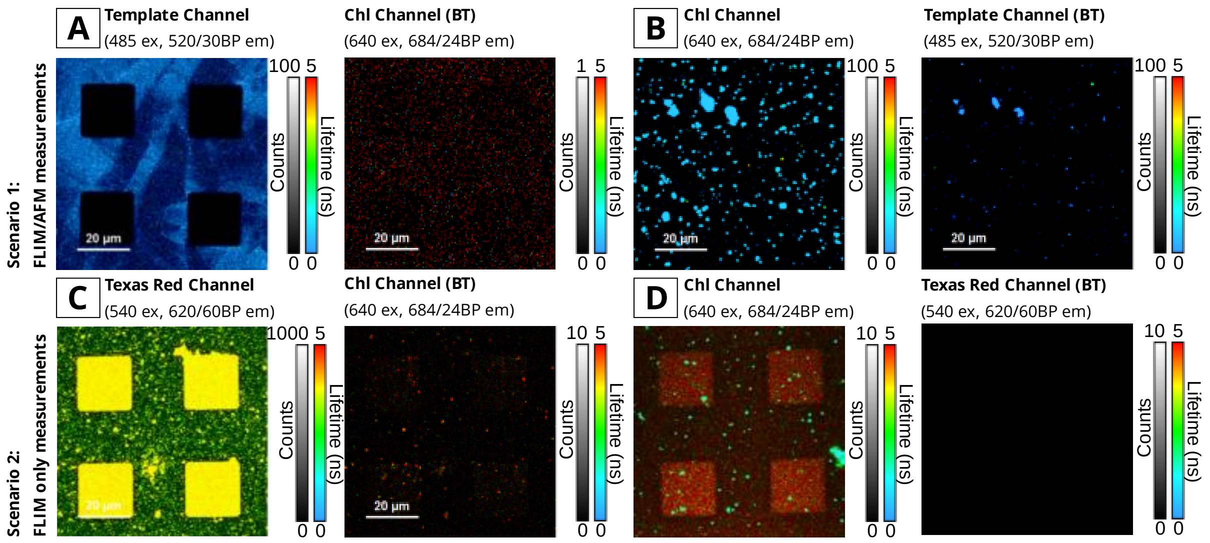


### Figure S2

Examples of the spectral crossover between FLIM channel for two scenarios. In the first scenario, for correlated FLIM+AFM measurements, we quantify the spillover between the intrinsic polymerized lipid fluorescence and chlorophyll-related channels. **(A)** FLIM image of a polymerized lipid sample (photo-patterned Diyne-PC), as observed in the “Template channel” (excited at 485 nm, emission at 520 nm with a bandpass width of 30 nm), and the spillover (BT) into the “Chl channel” (excitation at 640 nm, emission at 684nm with a bandpass width of 24 nm). The spectral spillover of the polymerized lipid emission to the Chl channel is less than 1%. **(B)** FLIM image of a sample containing non-ruptured thylakoid extracts, as observed in the Chl channel and the Template channel. The chlorophyll pigments can be excited in their Soret band, which overlaps significantly (2.78%) with the Template channel. However, since the polymerized lipid is acting as a pattern scaffolding, and not the focus of this study, we believe that this overlap into the Template channel is acceptable. In the second scenario, FLIM only measurements are optimized for the detection of Chl fluorescence and Texas Red fluorescence. **(C)** FLIM image of a patterned lipid bilayer composed of 99.5% DOPC + 0.5% TR DHPE in the “Texas Red channel” (excitation at 540 nm, emission at 620 nm with a bandpass width of 60 nm), and the Chl channel (optics as above). The spectral spillover from the TR emission to the Chl channel is less than 1% **(D)** FLIM image of hybrid thylakoid membranes (containing no TR) as seen in both the Chl channel, and spillover into the Texas Red channel. The spectral overlap from the chlorophyll emission into the Texas Red channel is 10.5 ± 1.7 %, however since the TR signal is typically 50-100 times higher than the Chl signal, the overall contribution is negligible.

FLIM measurements were acquired in two scenarios, for which the fluorescence spillover was quantified between the fluorescence channels of interest.

*Firstly,* correlated FLIM-AFM measurements were used to probe the topography of thylakoid/DOPC hybrid membranes as they backfilled into template polymerized lipid corrals (Figures 2, 3 and 4 in the main text). In order to be confident about the detected fluorescence signal, we quantified the fluorescence spillover on samples containing polymerized lipids only (“empty” scaffold patterns) and extracted thylakoids only (extracted thylakoids adhered to glass). Multiple fields of view were acquired per sample, using the same laser settings (pulse rate, intensity, etc) as would be used for subsequent measurements. The average number of fluorescence counts for the control samples are shown below in Table S1, where the spill over between the two channels has been calculated. In each case, the spill over between the two fluorescent channels was found to be negligible.

| **Sample** | **Template channel** | **Chl channel** | **Spill over to Template** | **Spill over to Chl** |
|---|---|---|---|---|
| # | Cnts | Cnts | % | % |
| Polymerized lipid template | $3.58 \pm 0.46 \times 10^6$ | $5.09 \pm 1.96 \times 10^3$ | N/A (100) | $0.12 \pm 0.01$ |
| Isolated thylakoids | $1.22 \pm 0.57 \times 10^6$ | $4.41 \pm 0.68 \times 10^7$ | $2.78 \pm 0.42$ | N/A (100) |

### Table S1

Mean counts from control samples in order to calculate the extent of channel spill over from the polymerized lipid channel (excited at 485nm, emission detected at 610-630 nm) to the Chl FLIM channel (excited at 640 nm, emission detected from 672-696 nm) and vice versa.

*In the second scenario,* standalone FLIM measurements were used to quantify spill over when the fluorophore TR was introduce into the sample to track lipid deposition. In this scenario, TR was excited using a 560 nm laser pulse, and detected in the range from 590-650 nm (TR channel). The Chl fluorescence was excited using a 640 nm laser pulse and detected in the range from 672 – 696 nm (Chl channel). FLIM measurements of hybrid membranes (No TR) and TR-only (TR-DOPC) control samples show that there is minimal overlap from Chl emission into the TR channel and vice versa. Table S2, below, shows the mean number of counts inside the corrals (N = 16 corrals) for various samples in each FLIM channel. For the hybrid membranes sample (Table S2, row 1), the crossover from the Chl fluorescence to the TR channel was calculated to be 10.49 ± 1.69 %. However, number of counts in the TR channel is approximately 40-fold higher than that in the Chl channel, therefore the Chl crossover contributes less than half a percent of the total counts in the TR channel and is negligible. From the DOPC-TR sample (Table S2, row 2), the crossover from the TR to the Chl channels was calculated to be 0.07 ± 0.02 %, Therefore in a hybrid membranes sample also containing TR, after considering the relative magnitudes of the TR and Chl signals, the crossover from the TR fluorescence is expected to account for ~3% of the total counts in the Chl channel. In either case, the crossover between Chl and TR channels is sufficiently small, that we can be confident in the fluorescence lifetimes measured in these channels.

| **Sample**<br>**(a) hybrid membranes**<br>**(b) lipid membranes** | **Chl channel** | **TR channel** | **Spill over to Chl** | **Spill over to TR** |
|---|---|---|---|---|
| # | Cnts | Cnts | % | % |
| Thylakoids/DOPC (a) | $1.08 \pm 0.08 \times 10^6$ | $1.14 \pm 0.13 \times 10^5$ | N/A (100) | $10.49 \pm 1.69$ |
| DOPC +Texas Red (b) | $3.09 \pm 0.57 \times 10^4$ | $4.41 \pm 0.68 \times 10^7$ | $0.07 \pm 0.02$ | N/A (100) |

### Table S2

Mean counts observed inside the corrals (N=16 corrals) of either hybrid membranes (thylakoids/DOPC) or lipid membranes containing 0.5% TR lipids. The extent of fluorescence spill over is calculated for both channels.

#### 3. Example FLIM images of empty polymerized lipid templates

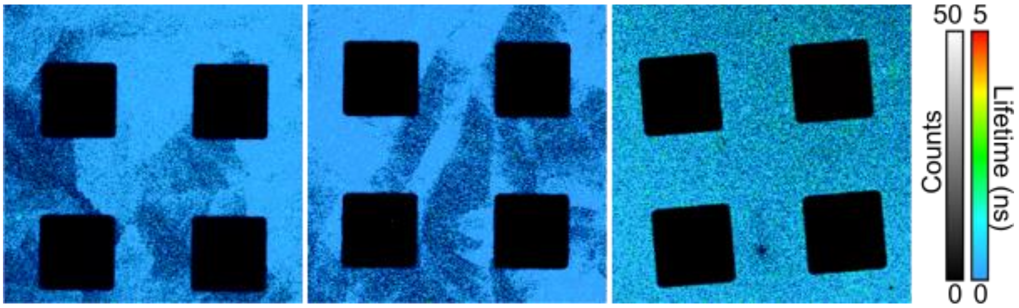


#### Figure S3

Multiple FLIM images of empty polymerised templates. Diyne-PC SLBs were polymerized by exposure to UV light through a photomask. Non-polymerized regions are then washed away by detergents, resulting in a patterned SLB with empty spaces and high-energy exposed bilayer edges.

#### 4. Hybrid membranes are structurally consistent across multiple preparations – FLIM gallery and homogeneity analysis

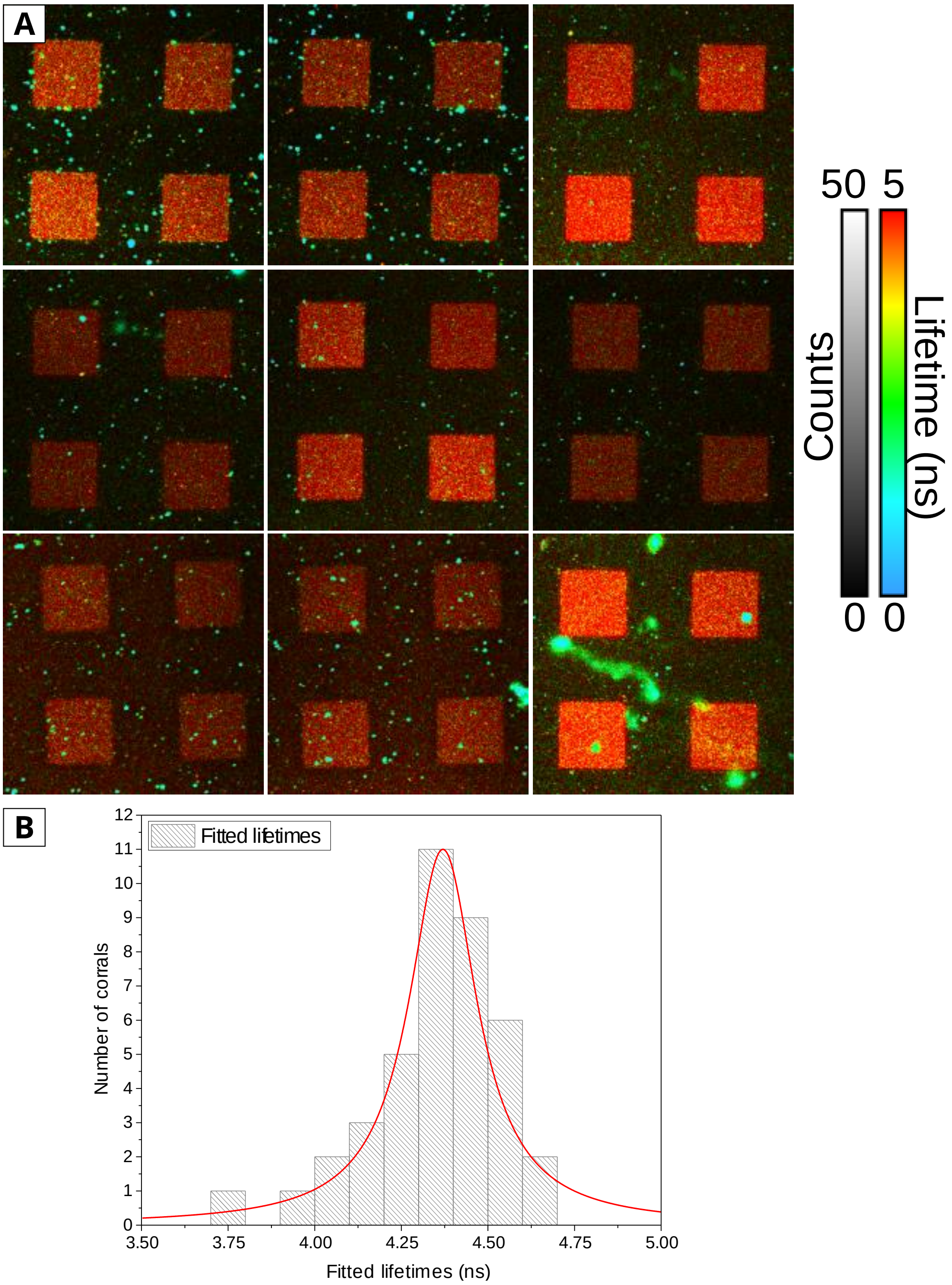


### Figure S4

**(A)** Multiple FLIM images of hybrid membranes within polymerized Diyne-PC templates. Samples were prepared across multiple different days and are seen to be highly reproducible in terms of their array patterned structure. The fluorescence intensity observed did vary between corrals (some are clearly brighter than others) but this could be due to a degree of photobleaching or other effects. Importantly fluorescence lifetime is similar seen here as a similar red color. **(B)** A histogram of fitted fluorescence lifetimes accumulated from 40 individual corrals. The fluorescence lifetime is very similar between hybrid membrane samples with a narrow distribution of Chl lifetimes (FWHM = 0.12 ns). This suggests the protein-protein interactions within the membrane are similar across multiple preparations.

#### 5. FLIM gallery of LHCII proteoliposomes adhered onto glass and calculations from these images to estimate the average fluorescence intensity per proteoliposome

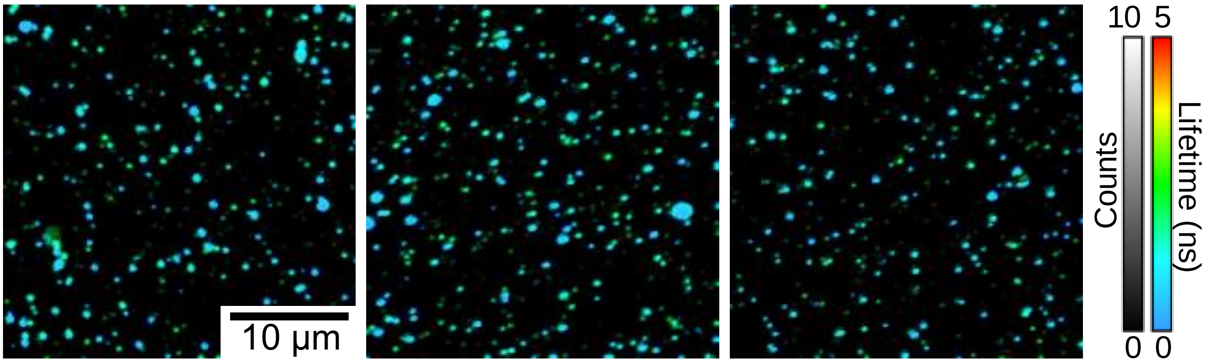


#### Figure S5

Multiple fields of view of LHCII proteoliposomes (containing approximately 2.8 μM LHCII and 1 mM thylakoid lipids) adhered onto cleaned glass. The samples were imaged in a buffer containing 40 mM NaCl, 20 mM, HEPES, pH 7.5 using the same acquisition settings and laser fluence used for the hybrid membranes, giving a direct comparison of the fluorescence intensity of a known amount of LHCII.

| **Proteoliposome number** | **Frames** | **Noise per pixel per frame** | **Number of selected pixels** | **Total counts** | **Detector noise in the selected area** | **Corrected $F_{vesicle}$** |
|---|---|---|---|---|---|---|
| | | Counts | Px | Counts | Counts | Counts |
| 1 | 500 | $3.68 \times 10^{-4}$ | 42 | 155 | 7.73 | 147.27 |
| 2 | 500 | $3.68 \times 10^{-4}$ | 32 | 17 | 5.89 | 11.11 |
| 3 | 500 | $3.68 \times 10^{-4}$ | 21 | 71 | 3.86 | 67.14 |
| 4 | 500 | $3.68 \times 10^{-4}$ | 39 | 30 | 7.18 | 22.82 |
| 5 | 500 | $3.68 \times 10^{-4}$ | 30 | 47 | 5.52 | 41.48 |
| 6 | 500 | $3.68 \times 10^{-4}$ | 20 | 71 | 3.68 | 67.32 |
| 7 | 500 | $3.68 \times 10^{-4}$ | 23 | 126 | 4.23 | 121.77 |
| 8 | 500 | $3.68 \times 10^{-4}$ | 34 | 182 | 6.26 | 175.74 |
| 9 | 500 | $3.68 \times 10^{-4}$ | 63 | 18 | 11.59 | 6.41 |
| 10 | 500 | $3.68 \times 10^{-4}$ | 57 | 152 | 10.49 | 141.51 |

#### Table S3

Calculation of the fluorescence intesnity signal for 10 example LHCII proteoliposomes (from images as shown in Figure S5). This "corrected" signal represents the number of counts (photons emitted) per proteoliposome after subtracting the detector noise. For a population of N=100 proteoliposomes, the average fluorescence intensity was found to be 74.58 ± 7.66 counts for a 500-frame acquisition. This equates to an **average $F_{vesicle}$** of 0.149 counts per frame.

### 6. Calculations for the average number of proteins within typical LHCII proteoliposomes

| Scenario | vesicle diameter, D (nm) | L/P (mol/mol) | $A_{vesicle}$ (nm$^2$) | $A_{lipid}$ (nm$^2$) | $A_{LHCII}$ (nm$^2$) | n (LHCII /vesicle) | density (LHCII /µm$^2$) |
|---|---|---|---|---|---|---|---|
| best estimate | 60 | 1071 | 11310 | 0.5 | 50.3 | 35.5 | 3143 |
| max. estimate | 100 | 1071 | 31416 | 0.45 | 38.5 | 21.0 | 2680 |
| min. estimate | 50 | 1071 | 7854 | 0.55 | 78.5 | 112.4 | 3577 |

### Table S4

Calculations for the number of LHCII proteins found on average per proteoliposome, given the estimated dimensions for the protein, lipids and the vesicle. This is for the proteoliposome sample shown in Figure S5 and Table S3 (which has 2.8 µM LHCII and 1 mM thylakoid lipids).

**D**, average diameter measured via dynamic light scatterting measurements (DLS), 60 nm is the average but given the accuracy of DLS the low and high values shown represent reasonable low and high estimates;

**L/P**, the average lipid-to-LHCII trimer ratio, as determinated from ensemble absorption spectroscopy measurements and spectral decomposition analysis using published methodology;

**$A_{vesicle}$**, calculated from $4\pi r^2$ (where, r= D/2);

**$A_{lipid}$**, published value for DOPC headgroup area,[8] given the uncertainty we use 0.45 and 0.55 as the low and high estimates;

**$A_{LHCII}$**, estimation of the membrane area occupied by one LHCII, from the consideration of space-filling models of published protein structures[8, 9] and then approximation of LHCII as a circular area ($\pi r^2$) where r = 3.5, 4.0 or 5.0 for the low, medium and high estimates (range due to uncertain protein packing);

Area per vesicle approximates to the following equation (note, the factor 0.5 is due to 2 lipids one from each two monolayer togethe to form one bilayer and thus occuping an area of $A_{lipid}$):

$$A_{vesicle} = n[A_{LHCII} + 0.5(L/P)A_{lipid}]$$

This expression was solved to calculate **n** using the values for **L/P**, **$A_{vesicle}$**, **$A_{lipid}$** and **$A_{LHCII}$**.

**Density** = n / $A_{vesicle}$.

The minimum, maximum and best estimates are made using the different possible values shown for each term, as shown.

#### 7. Calculations for the average number of proteins within a corralled hybrid membrane

| Scenario | $N_{LHCII/vesicle}$ | Measured average $F_{vesicle}$ | Measured $\tau_{vesicle}$ | Calculated $F_{LHCII}$ | Measured $F_{corral}$ | $N_{LHCII/corral}$ | Density (LHCII /corral) | $A_{LHCII}$ (area/ LHCII ) | $A_{protein}$ (%) |
|---|---|---|---|---|---|---|---|---|---|
| | # | cnts/frame | ns | cnts/frame | cnts/frame | #/ corral | #/ μm² | nm² | % |
| best estimate | 35.5 | 0.149 | 0.637 | 0.0263 | 1503 | 57100 | 143 | 50.3 | 0.72 |
| min. estimate | 21.0 | 0.149 | 0.637 | 0.0446 | 1503 | 33700 | 84 | 38.5 | 0.32 |
| max. estimate | 112.4 | 0.149 | 0.637 | 0.0083 | 1503 | 180000 | 451 | 78.5 | 3.54 |

### Table S5

Calculations for the number of proteins per corral, in terms of "LHCII-equivalents". This uses the average fluorescence counts measured by FLIM for an LHCII proteoliposome and converts to counts per LHCII protein, given the measured number of proteins within a typical proteoliposome (from Table S4). Consistent acquisition parameters were used to record FLIM images of both LHCII proteoliposomes and hybrid membranes.

**$N_{LHCII/vesicle}$**, estimated number of LHCII-equivalents per proteoliposome (**n** from Table S4). This range from the minimum to the maximum considering our combined uncertainties;

**$F_{vesicle}$**, estimated fluorescence intensity per proteoliposome per frame (**average $F_{vesicle}$** as caluclated in the caption of Table S3);

**$\tau_{vesicle}$**, the measured mean fluorescence lifetime of a typical LHCII proteoliposme (mean of N=100 measured particles from images similar to Figure S5);

**$F_{LHCII}$**, the FLIM counts expected per LHCII per frame calculated for each possible **$N_{LHCII/vesicle}$**, as follows. LHCII within proteoliposomes is known to self-quench, shortening the fluorescence lifetime due to the self-association of neighbouring LHCII.[2] The measured $\tau_{vesicle}$ of proteoliposomes of 0.637 ns (SD = 0.015 ns) implies significant quenching relative to isolated LHCII in detergent ($\tau_{DDM} \approx$ 4ns), so to crudely take this into account we can multiply by the ratio of the lifetimes (4/0.637). Thus, the intensity of the proteins in the unquenched state is estimated as:

$$\mathbf{F_{LHCII}} = \frac{\mathbf{F_{vesicle}}}{\mathbf{N}_{\left(\frac{\mathbf{LHCII}}{\mathbf{vesicle}}\right)}} \times \left(\frac{\tau_{DDM}}{\tau_{vesicle}}\right) = \frac{0.149}{\mathbf{N}_{\left(\frac{\mathbf{LHCII}}{\mathbf{vesicle}}\right)}} \times \left(\frac{4}{0.637}\right) = \frac{0.94}{\mathbf{N}_{\left(\frac{\mathbf{LHCII}}{\mathbf{vesicle}}\right)}}$$

**$F_{corral}$**, average fluorescence intensity measured in FLIM of hybrid membranes, as total counts within one corral per frame. This vale is found from careful analysis of the corrals from many images of hybrid membranes similar to those shown in Figure S4 (N = 16 corrals);

**$N_{LHCII/corral}$**, is the estimated number of LHCII trimers per corral, $N = F_{corral} / F_{LHCII}$;

**Density (LHCII/corral)** = **$N_{LHCII/corral}$** / $A_{corral}$ (where each corral has area $A_{corral}$ = 400 μm²);

**$A_{LHCII}$**, is the area occupied by a single trimeric LHCII protein complex, as estimated in Table S4;

**$A_{protein}$(%)**, estimated surface area fraction of the corral occupied by LHC and PS proteins:

$$A_{protein}(\%) = Density(\mu m^2) \times A_{LHCII}(nm^2)/10^6$$

### 8. Derivation of the Langmuir isotherm model

In the Langmuir isotherm model, the rate of material deposition is proportional to the amount of remaining free space on the surface. This assumes that each absorption site is energetically identical.[10]

This can be expressed mathematically as $\frac{dn}{dt} = R\left(1 - \frac{n}{N}\right)$, where n is the number of absorped molecules, R is a constant that describes the rate of absorption, and N is the total number of possible sites.

To solve this differential equation, we can consider two possible scenarios. In the first scenario, $n \ll N$, and there are a large number of remaining empty sites. The surface is not close to saturation.

In this scenario,

$$\frac{dn}{dt} = R\left(1 - \frac{n}{N}\right) \approx R$$

and therefore, $n = Rt + C$, where C is a constant arising from the integration of $\frac{dn}{dt}$. The rate of deposition is initially linear. This scenario was applied to fit the deposition of extracted thylakoids, where the surface coverage was very low throughout the incubation period.

In the second scenario, $n \sim N$, and the rate of deposition slows down as the number of free sites decreases.

$$\frac{dn}{dt} = R\left(1 - \frac{n}{N}\right)$$

which is rearranged to,

$$\frac{1}{1 - \frac{n}{N}}\, dn = R\, dt$$

This can be integrated to

$$-Nlog(N - n) = Rt + C$$

and rearranged in to the final form, $n = N - e^{\left(-\frac{R}{N}t + C\right)}$, used to fit the data from both the hybrid membrane and DOPC-TR deposition curves.

#### 9. A comparison of lipid deposition rate to protein deposition rate in a sample containing fluorescently-tagged lipids

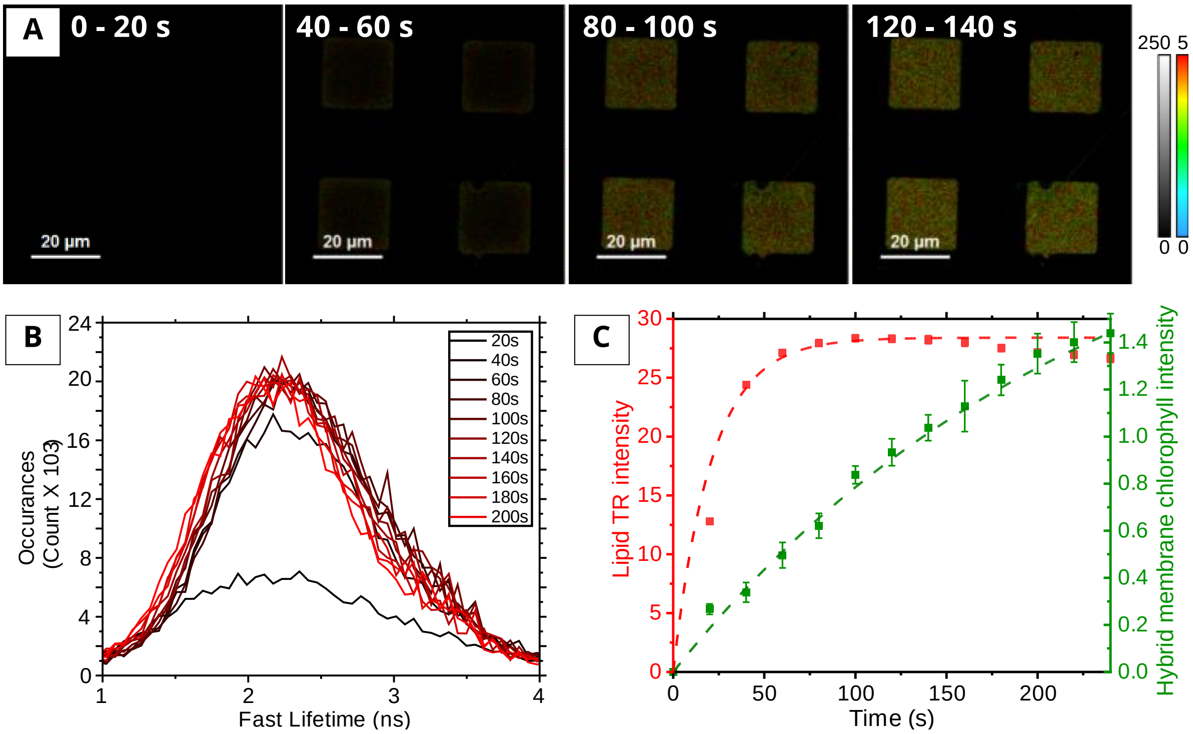


#### Figure S6

Analysis of the dynamic process of hybrid membrane formation. **(A)** A series of FLIM images showing the deposition of a hybrid membrane containing both photosynthetic proteins and fluorescently tagged (TR-DHPE) lipids. The FLIM channel shown here is optimized for the detection of Texas Red, in order to compare the rate of lipid deposition to the deposition of photosynthetic proteins (excitation at 561 nm and emission collected between 590-650 nm). **(B)** Evolution of the frequency distribution of lifetimes during the deposition. Dark red to light red represents the cumulative frequency distribution at increasing time points of 20-40s, 40-60s, 60-80s, 80-100s, 100-120s, 120-140s, 140-160s, 160-180s, 180-200s. Each distribution is fitted to a Gaussian curve, in order to calculate the Peak amplitude at each point. **(C)** Analysis of the growth over time of the TR intensity (from the peak amplitude data from panel B), compared to the hybrid membrane Chl intensity (from the long-lifetime Chl peak amplitude data from main text Figure 3). The lipid intensity saturates a lot sooner (~60s) than the hybrid membrane Chl intensity (~500s).

#### 10. Fluorescence Recovery After Photobleaching (FRAP) measurements to assess lipid and protein mobility in hybrid membranes

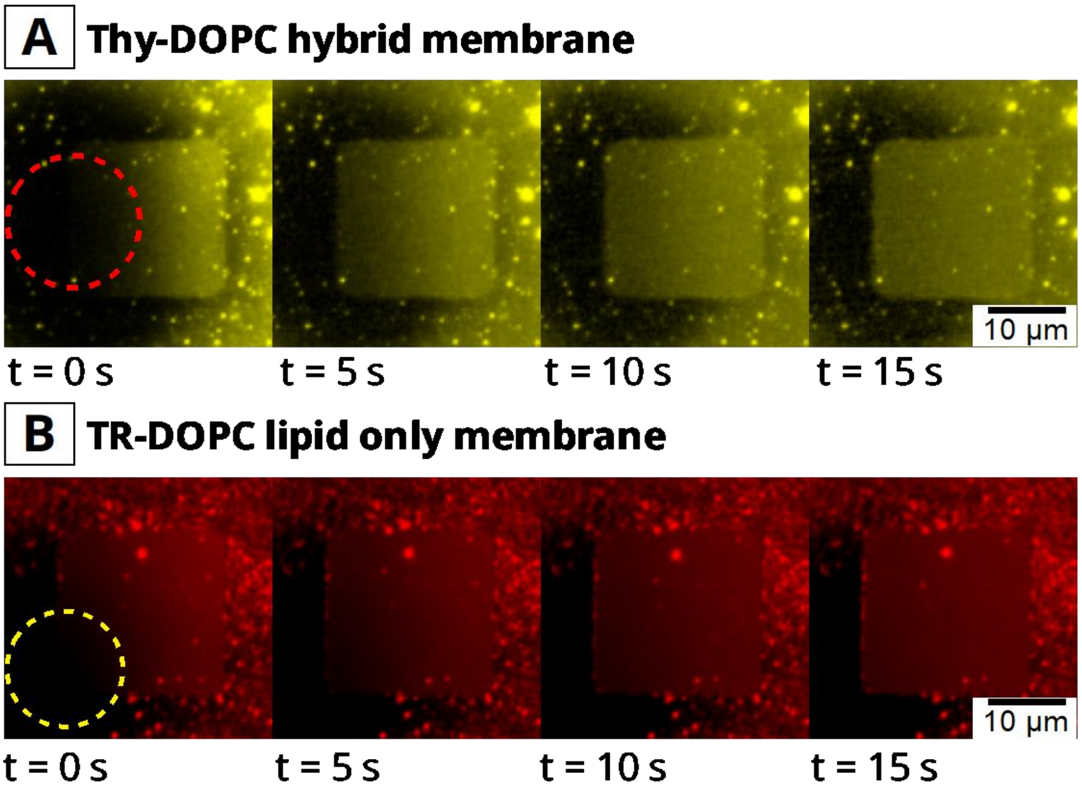


#### Figure S7

Epifluorescence FRAP experiments demonstrating protein and lipid mobility in hybrid membranes. **(A)** An epifluorescence time-lapse image series of hybrid membranes. Photosynthetic proteins are bleached within the *red* dashed region at t = 0 s, and non-bleached proteins can be seen to diffuse into the bleached region over the subsequent images. **(B)** An epifluorescence time-lapse image series of a patterned DOPC lipid membrane which contained 1% Texas Red DHPE fluorescently labelled lipids (mol/mol). The Texas Red fluorophores are bleached within the *yellow* dashed region at t = 0 s. Over the next 15s, the non-bleached lipids diffuse into the bleached area, until the fluorescence has fully recovered. The successful recovery of both protein and lipid fluorescence demonstrate the low levels of interaction between molecules and the substrate, and the free diffusion and connectivity of the membrane.

The epifluorescence instrument was a Nikon E600 microscope equipped with an Andor Zyla 4.2 sCMOS detector, set up with a filter cube optimized for Chl in for panel A (excitation 450-475 nm, dichroic 500 nm, emission 650-800 nm) and with a filter cube optimized for Texas Red in panel B (excitation 540-580 nm, dichroic 595 nm, emission 600-660 nm). Images were acquired using an x100 oil objective, 0.5s exposure and appropriate ND filters. For deliberate photo-bleaching, an aperture was inserted to expose circular region of ~15 μm diameter of the sample for a continuously period of 30 s at full power (i.e. no ND filters). Immediately following bleaching, full-field images were acquired sequentially in 5 second intervals.

### 11. Further data: AFM gallery of thylakoid membranes

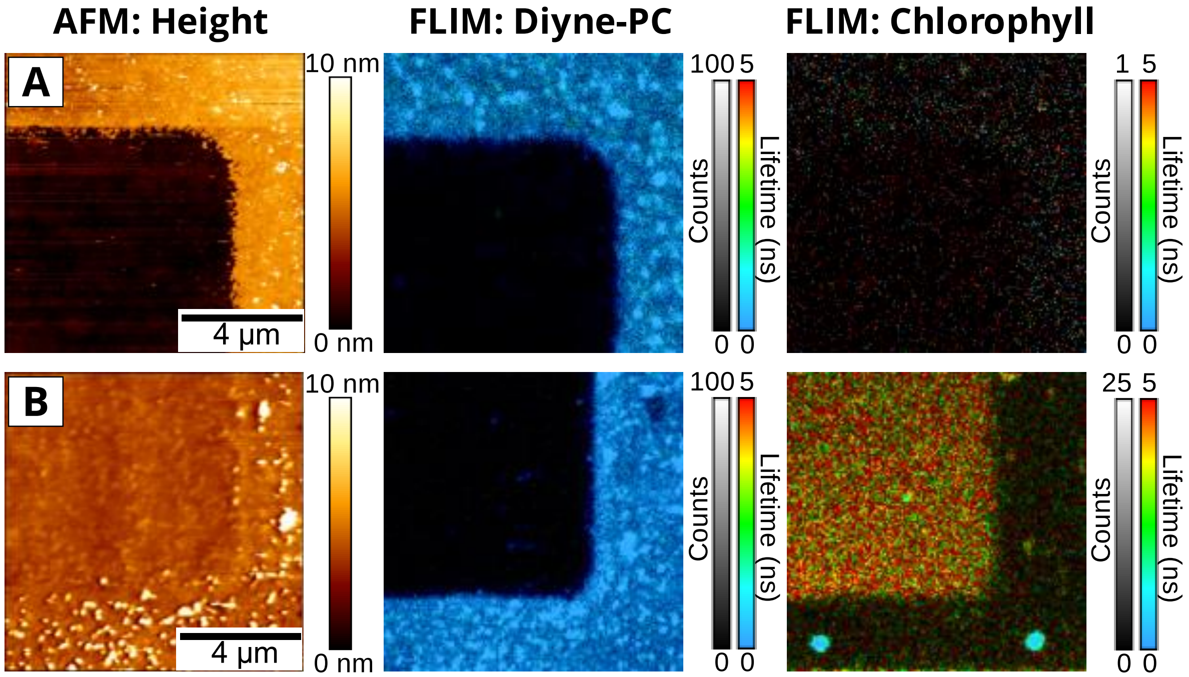


#### Figure S8

Additional correlated FLIM/AFM measurements at a higher magnification than those shown in the main text. For (A+B), left panel: AFM topograph; center panel: "polymer FLIM channel", i.e., optimized to detect the polymerized lipid fluorescence (laser excitation at 485 nm, emission bandpass between 505-535 nm); right panel: "chlorophyll FLIM channel", i.e., optimized to detect the fluorescence of chlorophylls which is expected from thylakoid LHC and PS proteins (laser excitation at 640 nm, emission bandpass between 672-696nm). **(A)** Correlated FLIM+AFM data showing a single square of the polymerized lipid "empty" template. The minimal signal in the chlorophyll FLIM channel is statistically indistinguishable from detector noise. **(B)** Correlated FLIM+AFM data showing a similar region as in (A), but after it was backfilled with the extracted thylakoids and liposomes to form the hybrid membrane within the corrals.

#### 12. Further data: AFM data of DOPC lipid bilayers backfilled into the polymerized lipid template

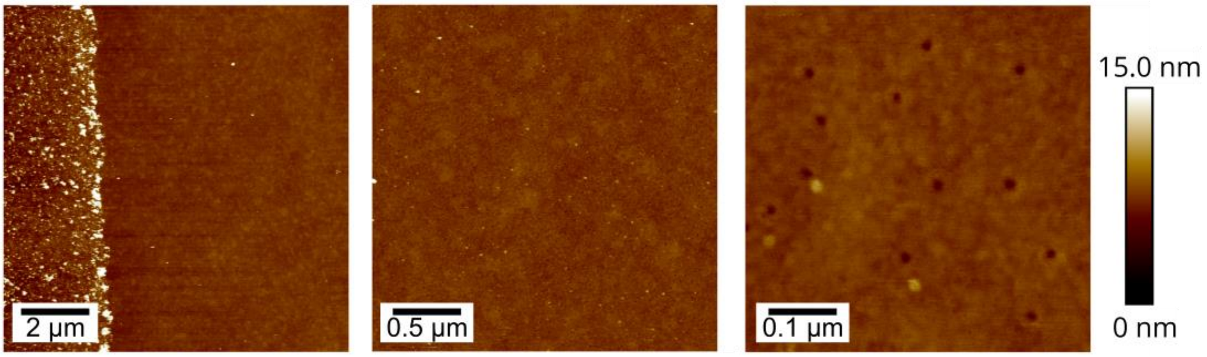


#### Figure S9

AFM measurements of showing that DOPC lipid membranes within a polymerized lipid template (sample prepared simply by backfilling a polymerized lipid template with DOPC lipid vesicles). The template edge can be seen on the left of the first AFM image. Inside the corral, the surface appears smooth and continuous with a low degree of roughness. This is in stark contrast to the >100nm wide and 4-5 nm deep pores observed in hybrid membranes. The only noticeable features are very small circular divots of approximately 1-3 nm in depth (see right-hand panel) which are commonly observed when forming SLBs on piranha-cleaned glass.[11]

#### 13. Classification of protein protrusion within defects in the bilayer

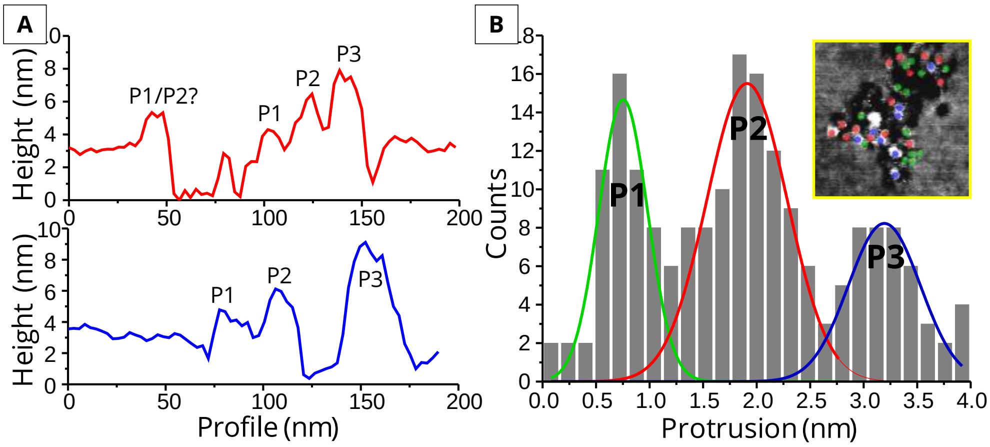


### Figure S10

Quantitative analysis of particle heights within a thylakoid hybrid membrane. **(A)** AFM height traces of particles within the defects, seem to reveal 3 distinct types of particles (labelled P1, P2 and P3). **(B)** A particle height histogram obtained from profile analysis across N=250 particles. Height traces were drawn across individual particles, as in (A), and the difference in particle height from the surrounding bilayer (protrusion) across many particles was accumulated as a histogram. Within this population, 3 peaks are observed with corresponding particle protrusions of 0.753 ± 0.027 nm (assigned P1), 1.911 ± 0.035 nm (assigned P2) and 3.194 ± 0.062 nm (assigned P3) respectively. The inset shows an example region, where particles have been color coded (P1 = *green*, P2 = *red*, P3 = *blue*) corresponding to their particle classification.

To classify the proteins, we used the much higher (~0.1 nm) z axis resolution of the AFM. Statistical analysis of the profiles across hundreds of proteins (N = 250) reveals a protrusion (i.e., the maximum height above the lipid bilayer) distribution with three distinct populations (see Figure 3I). These populations can be fitted to a Gaussian distribution ($R^2 > 0.99$), and were found to protrude from the bilayer by 0.753 ± 0.027 nm (assigned P1), 1.911 ± 0.035 nm (assigned P2) and 3.194 ± 0.062 nm (assigned P3) respectively. Our results are in moderate agreement with the known crystal structures of LHCII, PSII and Cytb$_6$f, and with previous AFM studies that have used the same approach.[12] The P1 peak is consistent with the predicted height for LHCII, and the P2 and P3 peaks show the same relative heights for PSII and Cytb$_6$f, respectively. From this we conclude that hybrid membranes contain many of the relevant photosynthetic membrane proteins.

#### 14. Further data: gallery of AFM images of hybrid membranes observed at a range of magnifications

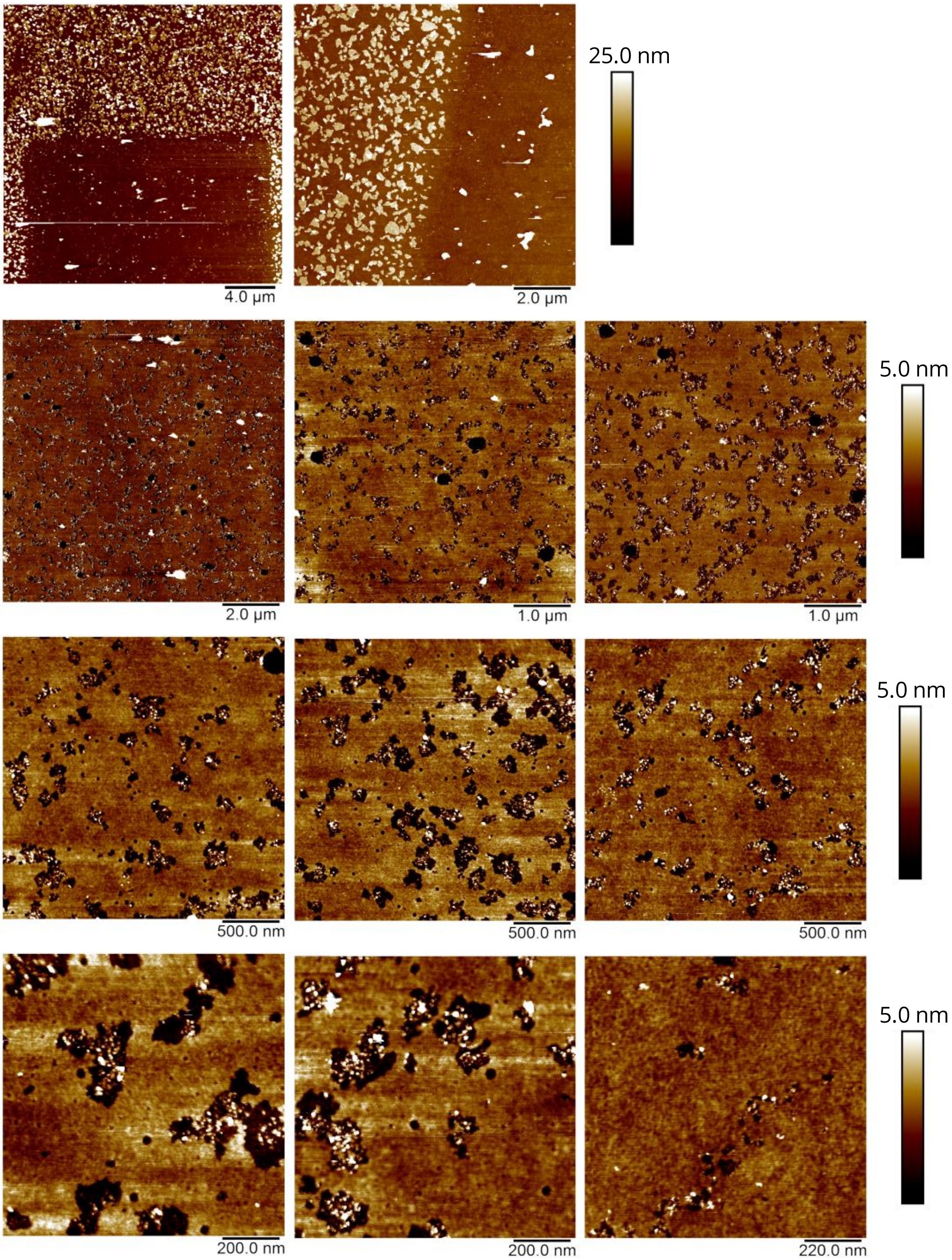


### Figure S11

Additional AFM height topographs of hybrid membranes (standard protocol of 1:3 thylakoids-to-DOPC backfilled into a polymerized lipid template). Scale bars are shown for each row of images.

#### 15. Additional spectroscopy data: lack of specificity in the photochemical assays

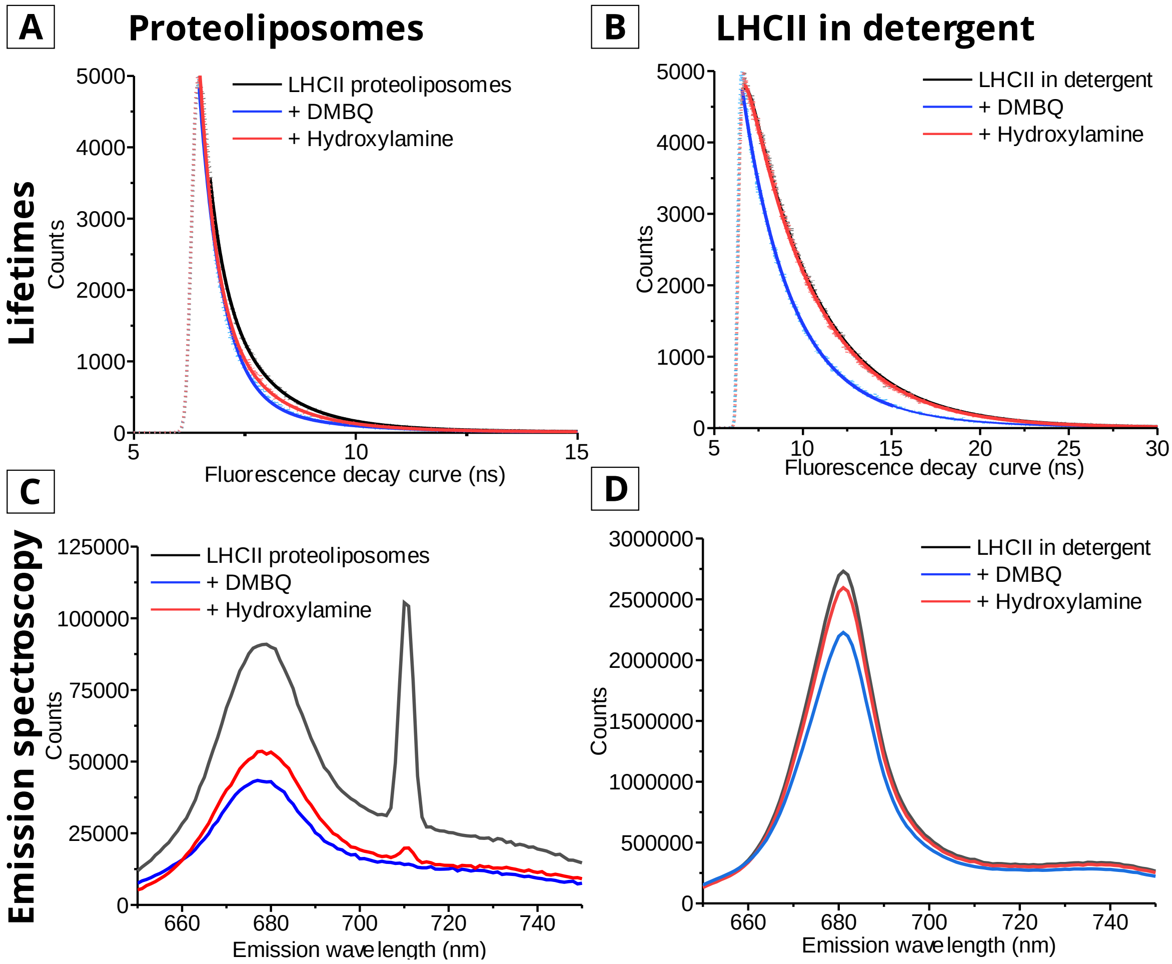


#### Figure S12

Cuvette-based steady-state and time-resolved fluorescence spectroscopy data of LHCII proteoliposomes (containing 10% w/w LHCII) and LHCII suspended in detergent micelles (in 0.3% α-DDM) in a buffer of 40 mM NaCl 20 mM HEPES pH 7.5. All measurements were performed in 10 x10 mm quartz cuvettes which were stirred with a magnetic stir bar to mix the solutions. **(A)** Fluorescence lifetime measurements of LHCII proteoliposomes, showing the decrease in fluorescence lifetime upon the addition of DMBQ, and the subsequent recovery of the fluorescence lifetime after the addition of hydroxylamine. **(B)** Fluorescence lifetime measurements of LHCII in detergent, showing the same effects of DMBQ and hydroxylamine as observed in proteoliposomes. **(C) & (D)** Emission spectroscopy plots for LHCII proteoliposomes and LHCII in detergent respectively. In both plots, there is a decrease in fluorescence emission after the addition of DMBQ, followed by the partial or full recovery after the addition of hydroxylamine. This evidence, combined with FLIM measurements (see figure 8) suggest that these photochemical assays, are not specific to the electron transfer photochemistry of PSII.

This cuvette-based fluorescence spectroscopy was performed using an Edinburgh Instruments FLS980 system equipped with dual excitation monochromators and dual emission monochromators. Samples were maintained at 20 °C and gently stirred using a thermoelectrically-cooled cuvette-holder with magnetic stirring. For steady-state emission spectra, excitation was at 473 nm and emission collected from 500-800 nm (2 nm and 1 nm bandwidth excitation and emission slits, respectively). Time-resolved fluorescence lifetime measurements used a 473 nm pulsed diode laser for excitation (0.5 MHz repetition rate and minimized power) and emission was collected at 681 nm with 10 nm bandwidth emission slits.

### 16. Control measurements to optimize FLIM acquisitions: selection of an excitation fluence which minimizes singlet-singlet annihilation artefacts

We wanted to minimize any risk of artefacts in fluorescence lifetime in FLIM measurements of photosynthetic proteins, which are known to occur if excitation power is set at too high a level.[2] [13] To confirm this, we varied the excitation power over a range expected to be above and below the level where singlet-singlet annihilation effects may occur. Figure S12 shows fitted lifetimes from FLIM data acquired for a range of photosynthetic samples (both extracted thylakoids and LHCII-only samples). For each sample, multiple different areas were imaged using excitation powers. The excitation power was modulated using a combination of ND filters and an adjustable microblade that obstructs part of the laser beam. The average power readout is provided in arbitrary units within in the SymPhoTime software, which is then converted into SI units using a linear power conversion relationship for each laser wavelength. The pulse FHWM was maintained at a minimal value (<90 ps) by setting the lasing voltage to the minimum threshold required to power the lasers. The pulse repetition rate was set to 10 MHz, such that the fluorophore could fully decay before the next pulse to avoid photon pile-up artefacts. For all lasers, we estimate the laser spot diameter to be ~800nm, allowing us to calculate the peak power per area (fluence) delivered to each sample.

The graph in Figure S12 shows how the average lifetime, <τ>, of each sample changes with the applied fluence (note: <τ> is on a logarithmic scale). In all samples, <τ> remains approximately constant at low fluences (below 0.01 mJ/cm$^2$), before exponentially decreasing above 0.01 mJ/cm$^2$ (the unacceptable fluence range is highlighted in red). In order to minimize the collection time required, whilst still maintaining a good signal and an acceptable signal to noise, we selected the upper end of the acceptable fluence range (0.01 mJ/cm$^2$) to be used for the FLIM acquisition of subsequent samples.

All the lasers share the same ND filter and microblade aperture, however, due to the differences in beam shape and beam diameter, the microblade cut off affects all three lasers differently. Therefore, when operating in Pulsed Interleaved Excitation mode (PIE) mode, setting the laser fluence for the photosynthetic excitation indirectly affected the laser power applied to the Texas Red change. The calculated laser fluences that we used in the current study are shown for 485/640 PIE (combined with the 485/640 dichroic mirror) and for 560/640 PIE (using the 561/640 dichroic mirror) in Table S6 and S7, respectively.

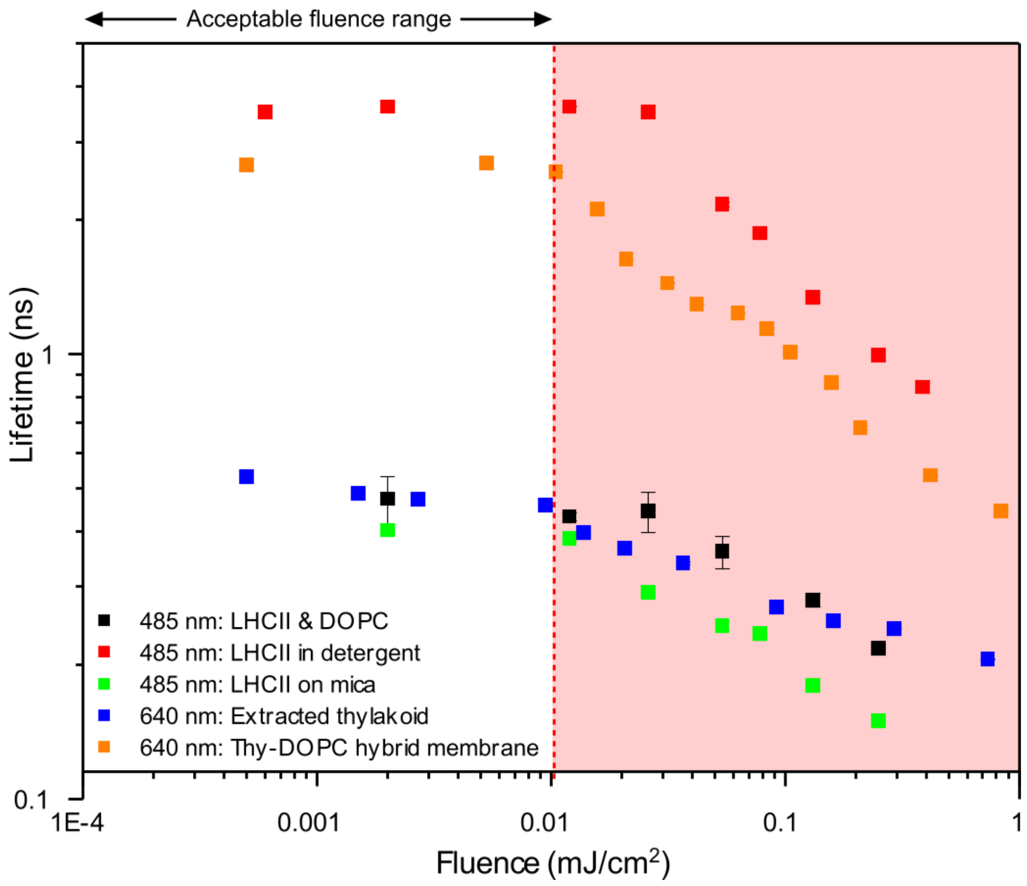


### Figure S13

Average fluorescence lifetime versus excitation fluence across multiple photosynthetic samples, highlighting the acceptable fluence range where lifetime artefacts are minimal.

| Wave-length | Energy per photon | Laser spot diameter | Rep rate | Pulse FWHM | Average laser power output | Average intensity | Peak power | Fluence | Fluence with obj. loss |
|---|---|---|---|---|---|---|---|---|---|
| nm | J | nm | MHz | ps | W | W/cm$^2$ | J/s | mJ/cm$^2$ | mJ/cm$^2$ |
| 485 | 4.10E-19 | 800 | 10 | 90 | 1.50E-6 | 2.98E2 | 1.67E-03 | 0.030 | 0.025 |
| 640 | 3.10E-19 | 800 | 10 | 70 | 6.85E-7 | 1.36E2 | 7.61E-04 | 0.014 | 0.012 |

### Table S6

Calculations for the average laser fluence for the FLIM operating in 485/640 PIE mode using the 485/640 dichroic mirror. The 640 nm laser pulse is specifically targeted for the chlorophyll fluorescence, and as such we chose to maintain a low fluence where singlet-single annihilation effects are minimal. This configuration was used when acquiring both the "Diyne-PC FLIM channel" and "Chlorophyll FLIM channel" (main text Figure 4, supplementary Figure S8).

| Wave-length | Energy per photon | Laser spot diameter | Rep rate | Pulse FWHM | Average laser power output | Average intensity | Peak power | Fluence | Fluence with obj. loss |
|---|---|---|---|---|---|---|---|---|---|
| nm | J | nm | MHz | ps | W | W/cm$^2$ | J/s | mJ/cm$^2$ | mJ/cm$^2$ |
| 560 | 3.55E-19 | 800 | 10 | 70 | 1.23E-6 | 2.45E2 | 1.76E-3 | 0.024 | 0.021 |
| 640 | 3.10E-19 | 800 | 10 | 90 | 6.85E-7 | 1.36E2 | 7.61E-04 | 0.014 | 0.012 |

### Table S7

Calculations for the average laser fluence for the FLIM operating in 560/640 PIE mode using the 560/640 dichroic mirror. The 640 nm laser pulse is specifically targeted for the chlorophyll fluorescence, and as such we chose to maintain a low fluence where singlet-single annihilation effects are minimal. This configuration was used when acquiring both the "Texas Red FLIM channel" and "Chlorophyll FLIM channel" (supplementary Figure S5).

#### 16. Confirmation of the validity of lifetime measurements made in the absence of an O2 scavenging enzyme

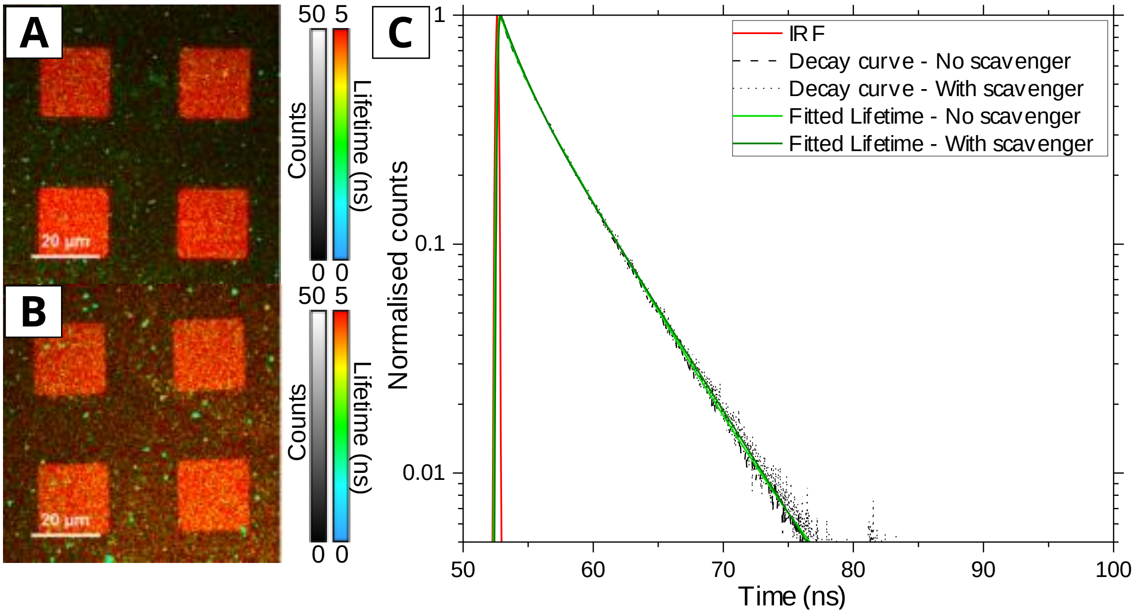


#### Figure S14

**(A)** Hybrid membrane that has been prepared and imaged as normal, as described in the main text. **(B)** A hybrid membrane prepared as usual, but subsequently washed into a buffer that has been sparged with dry $N_2$ gas and contains an $O_2$ scavenging enzyme system (2.5 mM protocatechuic acid and 50 nM protocatechuate-3,4-dioxygenase from *Pseudomonas* species).[14] After washing, the sample was allowed to stand for >1 hr at room temperature before imaging, to allow for full removal of any dissolved $O_2$ from the sample solution. **(C)** Comparison of fluorescence decay curves between the samples with or without the $O_2$ scavenger. After addition of the $O_2$ scavenger, there is no significant change in fluorescence lifetimes and only a minimal change in the fluorescence intensity (increased background on the template area), therefore, we can be confident that any dissolved $O_2$ in the prepared buffers has little effect on our hybrid membrane system.
