## Supplementary figures and images for "Understanding the photophysics and structural organization of photosynthetic proteins using model lipid membranes assembled from natural plant thylakoids"

### Graphic abstract

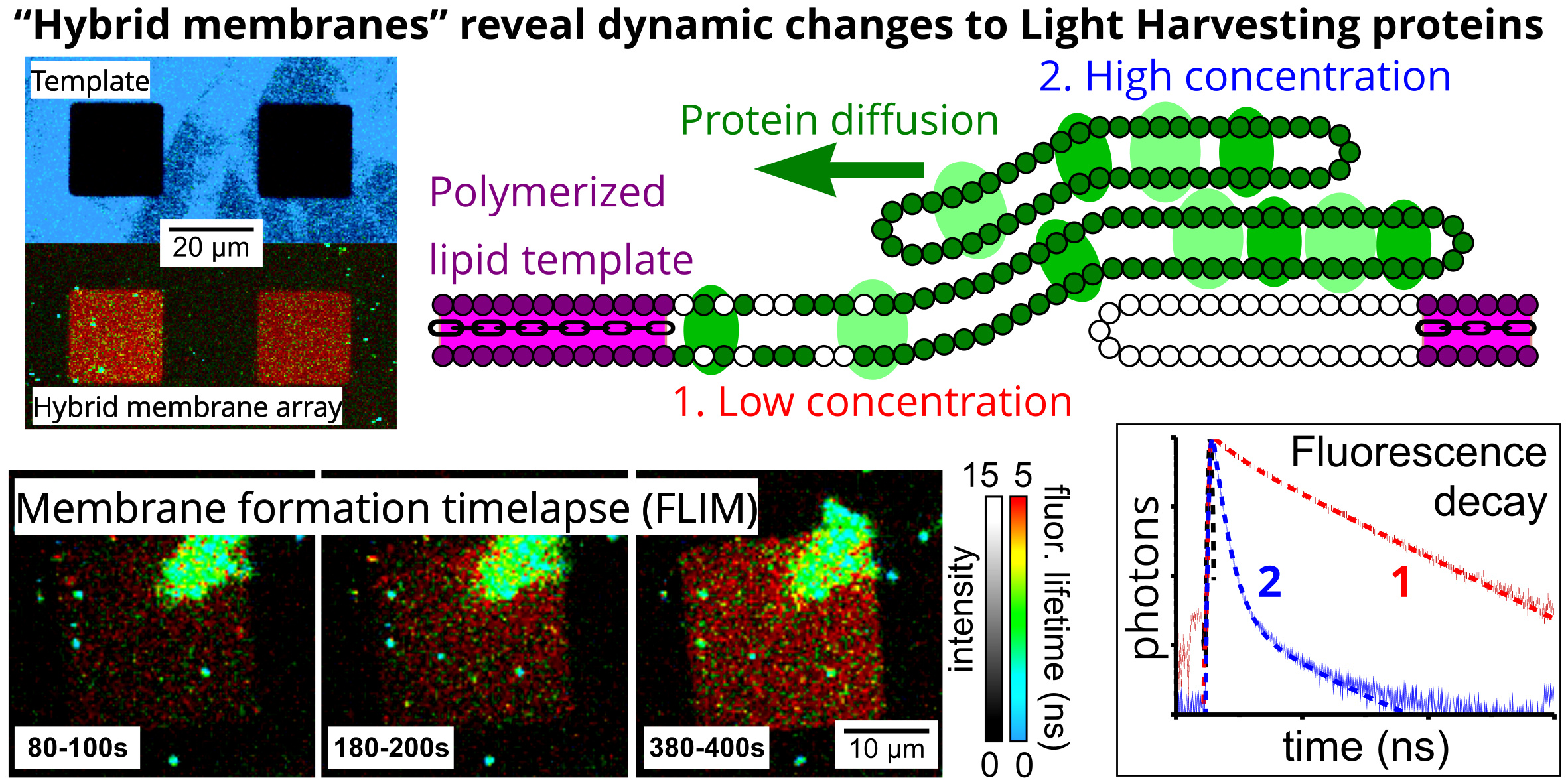
